# Asymmetric Hydration and Protonation Switching of Dual Aspartates Drive Flagellar Rotation

**DOI:** 10.64898/2026.04.14.718414

**Authors:** Jiancheng Luo, Haidai Hu, Zhitao Cai, Chen Shi, Yichong Lao, Peng Xiu, Nicholas M. I. Taylor, Yandong Huang, Yong Wang

## Abstract

The bacterial flagellar motor is an intricate nanomachine that transforms chemical energy from ion gradients into mechanical rotation, enabling bacterial movement. While stator unit architectures are conserved across species, the molecular link connecting ion translocation to rotational force generation remains elusive. In this study, we refined the cryo-EM structures of MotAB from *Campylobacter jejuni* (CjMotAB) and integrated a suite of approaches—including single-structure based p*K*_a_ predictors and free energy perturbation (FEP) calculations, as well as standard and constant-pH molecular dynamics (CpHMD) simulations of various structural models representing the plugged, unplugged, and plug-removed states with different protonation states of D22—to dissect its rotational mechanism. Based on p*K*_a_ calculations, the D22 residues in chains F and G of MotB were identified as proton carriers supporting the previous hypotheses. Importantly, we observed asymmetric hydration patterns of the two D22 residues in the MotB dimer, along with their hydrogen bonding interactions with MotA T189, which contribute to functional specialization. Our findings reveal that MotA rotation requires two essential prerequisites: plug removal and alternating D22 protonation switching, coupled with dynamic *gauche*-*trans* conformational changes in the sidechain of D22. This work clarifies how protonation dynamics and structural asymmetry synergistically regulate CjMotAB rotation, advancing our understanding of bacterial flagellar motor function and providing a foundational framework for investigating diverse ion-driven biological motors.

## 1 Introduction

In nature, the principles of adaptation and survival are fundamental. They drive the evolution of diverse traits across species. Many bacteria, to optimize nutrient acquisition and avoid harmful conditions, have developed specialized structures like flagella.^1,2^ Flagella are whip-like appendages that endow bacteria with motility. By rotating their flagella^3,4^ bacteria can move through dynamic environments, which significantly enhances their survival and adaptability.^5^

A bacterial flagellum is composed of three main components: the filament, the hook, and the motor. The motor, a highly complex nanomachine, consists of a large rotor and several smaller stator units.^6^ The rotor includes multiple substructures, such as the L ring, P ring, MS ring, C ring, the rod, and the export apparatus, which collectively transmit torque to the filament to drive bacterial movement.^7^ However, torque generation is powered by the stator units, which utilize cation concentration gradients to transform chemical energy into mechanical energy, thereby enabling flagellar rotation. These stator units are therefore critical for the functionality of the bacterial flagellum.^8^

The MotAB stator unit, predominantly found in motile Gram-negative bacteria, functions using a proton concentration gradient to power the bacterial flagellar motor.^9,10^ Interestingly, the recently identified ZorAB system, a widely distributed bacterial immune mechanism that defends against bacteriophage infections, has also been shown to utilize the proton gradient for rotational function.^11,12^ Beyond these proton-driven systems, other stator units’ mechanisms exhibit diverse ion preferences and roles. For example, the PomAB stator unit,^13–18^ found in *Vibrio* species, drives polar flagellum rotation using sodium ions, while MotPS,^19–23^ present in some bacteria, similarly relies on sodium ions for flagellar motility. In contrast, ExbBD, part of the Ton system in Gram-negative bacteria, harnesses the proton motive force to energize nutrient transport across the outer membrane, and TolQR uses the proton motive force to maintain outer membrane integrity.^24–28^ Despite these differences in ion utilization and biological function, structural biology studies reveal that all stator units—MotAB, ZorAB, PomAB, MotPS, ExbBD, and TolQR—share a highly conserved 5:2 rotary motor architecture.^29^ Remarkably, these bacterial stator units are homologous heteroheptamers, consisting of a homodimer of one subunit surrounded by a homopentamer of another subunit. While MotAB and ZorAB align in their use of protons, PomAB and MotPS diverge with sodium ion dependence, likely reflecting environmental adaptations. Functionally, MotAB, PomAB, and MotPS support motility, ZorAB aids in phage defense, and ExbBD and TolQR facilitate membrane processes, illustrating the stator unit architecture’s versatility across bacterial species.

It has been proposed that, upon exposure to external stimuli, the periplasmic peptidoglycan-binding (PGB) domain of the MotB dimer engages with the peptidoglycan layer and is pulled upward.^10,30^ This interaction is thought to trigger a conformational change in the stator unit, shifting it from a relaxed, inactive “plugged” state to an extended, active “unplugged” state. As a result, the ion channel opens, permitting the influx of protons (or other cations) down their electrochemical gradient. The resulting ion flow converts the stored chemical energy of the gradient into mechanical torque, which is transmitted to the C-ring of the rotor, thereby driving flagellar rotation (Fig. 1). Nevertheless, the precise molecular mechanism by which torque is generated remains far from being understood.

**Figure 1:**
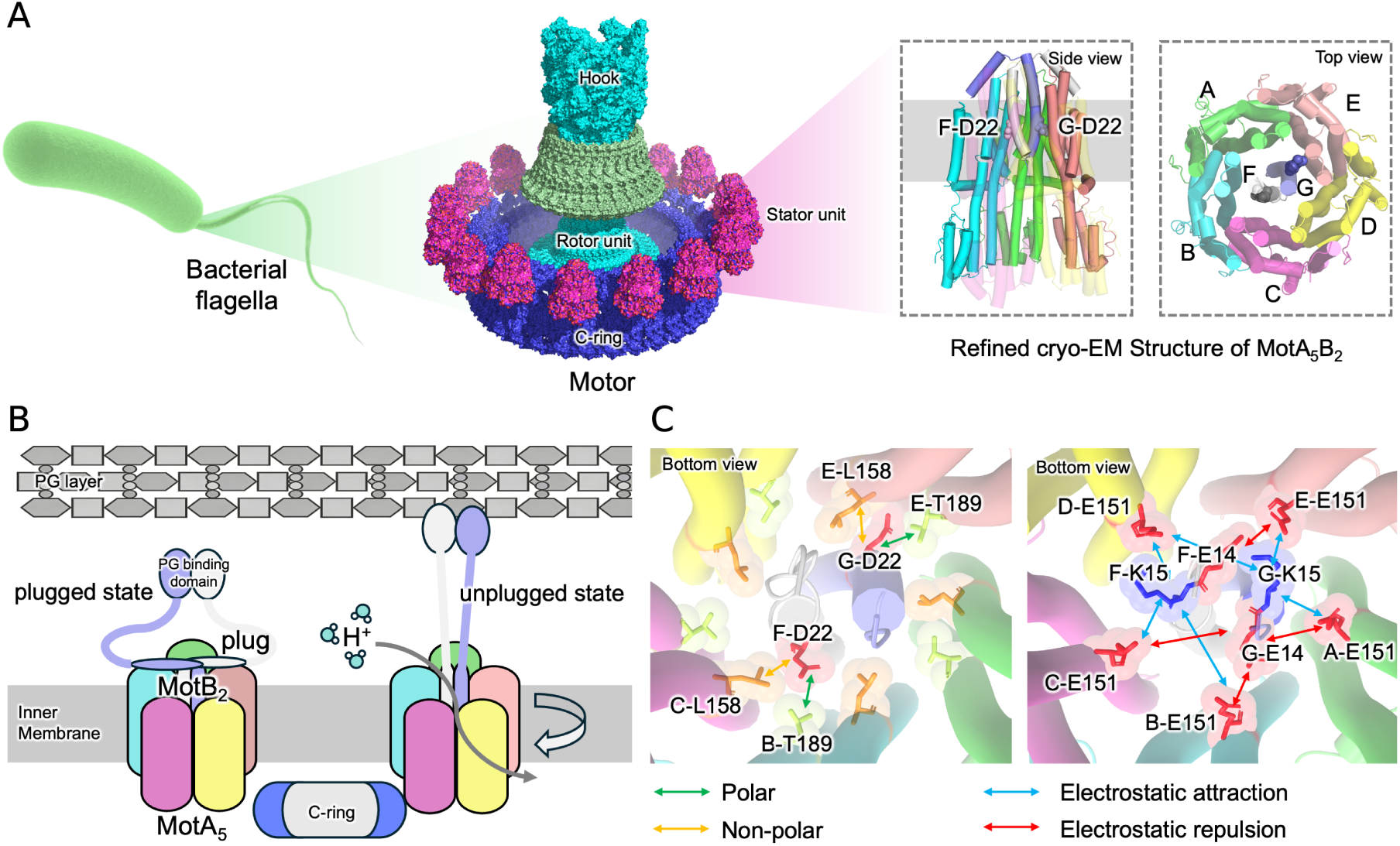
Structural and mechanistic overview of the bacterial flagellar motor CjMotAB. (A) Schematic hierarchical architecture of the bacterial flagellar motor system. Left: Schematic illustration of the flagellum–hook–motor assembly within a bacterial cell. Middle: Magnified view of the motor, emphasizing the rotor unit (teal/green) and membrane-embedded stator units (pink/purple). Right: High-resolution structure of the stator unit complex MotA_5_B_2_, depicting its heteromeric subunit organization. Chain A: green, chain B: cyan, chain C: magenta, chain D: yellow, chain E: salmon, chain F: gray, chain G: slate. (B) Mechanistic hypothesis of CjMotAB rotation: plugged vs. unplugged states. Depicts the MotAB stator unit complex in two functional states. In the plugged state (left), MotB localizes against the peptidoglycan layer, inhibiting ion flux. In the unplugged state (right), MotB undergoes ion channel opening, facilitating proton (H^+^) influx to drive MotA rotation. (C) Molecular interactions underpinning motor rotation visualized from the basal (bottom-up) perspective of CjMotAB. Diagrams illustrate force-generating interactions (polar, non-polar, electrostatic attraction/repulsion) between critical stator unit and rotor residues. Colored arrows denote interaction types (green = polar; yellow = non-polar; blue = electrostatic attraction; red = electrostatic repulsion), elucidating how these forces coordinate to drive motor rotation.

In this study, we focus on the *Campylobacter jejuni* CjMotAB stator unit complex. Previous single-particle cryo-electron microscopy (cryo-EM) studies have highlighted the critical role of the conserved aspartic acid residue at position 22 (D22) in the MotB dimer for stator unit function and motor rotation.^10^ This acidic residue is highly conserved across bacterial flagellar motor stator units. The unidirectional rotation of the bacterial flagellar motor’s MotAB stator units is thought to involve a dynamic interplay between protonation state transitions of MotB D22 and its interactions with hydrophobic (L158) and polar (T189) residues in the MotA pentamer.

More specifically, the transmembrane D22 residue functions as a regulated proton gate. In its deprotonated state (D22^D^), the negatively charged carboxylate attracts protons from the periplasm, leading to protonation and formation of the neutral state (D22^P^). Simultaneously, the hydrophobic leucine residue L158 in MotA helps stabilize the MotA–MotB interface via nonpolar interactions, preserving structural integrity during torque generation. Upon protonation, the newly formed D22^P^ establishes polar interactions with the threonine residue T189 of MotA. This MotA–MotB interaction is thought to induce a conformational change, resulting in an approximately 36° rotation of the MotA pentamer. This cyclic process—coupling proton translocation to mechanical rotation—likely relies on the hydrophobic scaffolding of L158 to maintain stator unit rigidity and the polar network of T189 to facilitate proton relay and conformational switching (Fig. 1). Collectively, these interactions are thought to convert proton-motive force into unidirectional rotation, highlighting the critical role of D22’s protonation state transitions in coordinating multi-subunit dynamics within the flagellar motor.

Besides, recent molecular dynamics (MD) simulations leveraging simplified coarse-grained (CG) structure-based models,^31^ where each residue is represented by a single *C_α_*bead,^32^ have been instrumental in unraveling key mechanistic insights into the bacterial flagellar motor. These CG MD simulations suggest that the glutamic acid residue at position 151 (E151) in MotA may act as a proton relay, accepting protons released by MotB D22 to play a critical role in the rotation mechanism. Notably, rotary motion was detected when three out of the five MotA E151 sites are protonated. Protonation of MotA E151 neutralizes its charge, disrupting its electrostatic attraction with the positively charged MotB K15. This disruption unlocks the conformation, enabling rotational dynamics driven by the electrostatic attraction between subsequent deprotonated MotA E151 sites and MotB K15 residues—an interaction that further generates the required torque.

However, the mechanistic hypothesis derived from these CG MD studies are subject to inherent limitations of the CG model, which oversimplifies side chain details and hydration effects. Furthermore, standard MD simulations (with fixed particle charges) cannot capture the dynamic protonation state transitions of residues that are central to the proposed mechanism.

To overcome these limitations, constant-pH molecular dynamics (CpHMD) methods have been developed, allowing protonation states to respond dynamically to conformational changes at a given pH. This approach allows for the simultaneous sampling of conformational and protonation states, facilitating the determination of protonation and tautomeric states of titratable residues—states that are experimentally challenging to characterize. Furthermore, p*K*_a_ values can be derived from CpHMD simulations across a range of pH conditions, allowing for conformational insights into the rotation mechanism.

In this work, we refined single-particle cryo-EM reconstructions of both plugged and unplugged CjMotAB to high resolution, resulting in structural models with improved geometric quality and completeness. Using these refined models, we performed explicit-solvent atomistic *λ*-dynamics CpHMD simulations implemented in GROMACS for membrane proteins.^33–37^ The integration of high-resolution structural data with CpHMD simulations enables more accurate characterization of protonation states and conformational dynamics of key residues involved in motor function.

For validation, we complemented CpHMD results with single-structure p*K*_a_ prediction methods (PropKa 3.0,^38^ PypKa,^39^ and DeepKa^40–42^) as well as free energy perturbation (FEP)-based p*K*_a_ calculations, which quantify residue-specific deprotonation free energy differences. By combining p*K*_a_ predictions from these approaches with CpHMD-derived protonation-coupled conformational dynamics, standard MD simulations of side-chain hydration and flexibility, and residue coevolution analyses, we propose a comprehensive proton-driven rotation model that extends and complements previous studies.

## 2 Methods

### 2.1 Cryo-EM Structure Refinement of CjMotAB

The two cryo-EM datasets of CjMotAB corresponding to its plugged and unplugged conformational states were reprocessed using cryoSPARC version 4.7.0.^43^ Unless specified otherwise, all data processing workflows followed the protocols described in a prior study.^16^ Patch motion correction was used to estimate and correct frame motion and sample deformation (local motion). Patch contrast transfer function (CTF) estimation was used to fit local CTF parameters to the micrographs. Micrographs were manually curated to remove poor-quality images (those with relative ice thickness greater than 1.1 and CTF values worse than 3.5 Å). Particles were picked using templates generated from the cryo-EM map from a previous study^10^ and extracted with a box size of 400 pixels. For both datasets, 2D classification was initially skipped, and 3D heterogeneous refinement was performed directly using one high-quality cryo-EM map from the previous study and two random maps. This step was iterated three times, retaining only particles from good classes. One round of 2D classification was then performed. Non-uniform refinement with a dynamic mask was applied to obtain a high-resolution map. Another round of non-uniform refinement was carried out with optimization of per-particle defocus and per-group CTF parameters. Reference-based motion correction was consecutively performed, followed by an additional round of non-uniform refinement and local refinement to further improve map resolution. Key experimental parameters for all datasets, including the number of curated micrographs, total electron exposure dose, number of particles used for final structural refinement, and the overall resolution of the resulting cryo-EM maps, are summarized in Table S1.

### 2.2 Theory and Implementation of CpHMD Simulation

CpHMD simulations were performed using a modified version of GROMACS 2021^35,44^ (available at https://gitlab.com/gromacs-constantph/constantph), with the *λ*-dynamics approach. In this continuous CpHMD method, *λ* represents the protonation state of a titratable residue, where *λ* = 0 corresponds to the protonated state and *λ* = 1 corresponds to the deprotonated state. The *λ* value evolves continuously between 0 and 1, governed by the following Hamiltonian equation:

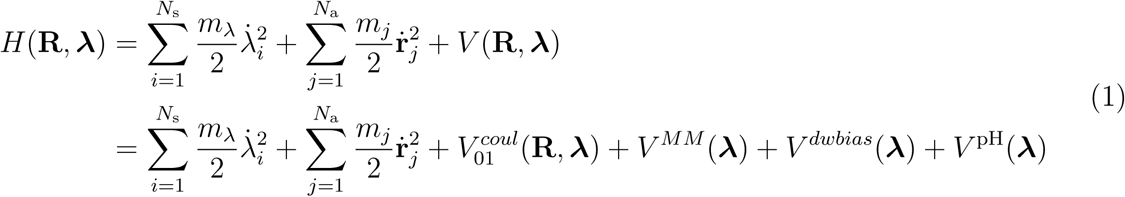

Here, i denotes a titratable site, ***λ*** is the vector of *λ_i_* coordinates for all *N_s_* titratable sites, **R** is the vector of Cartesian coordinates *r_j_* for all *N_a_* atoms with mass *m_j_*, and *m_λ_*is a fictitious mass of 5.0. The electrostatic potential *V_0_*_1_ *^coul^*(**R**, ***λ***) interpolates between protonated (*V_0_ ^coul^*) and deprotonated (*V_0_ ^coul^*) states:

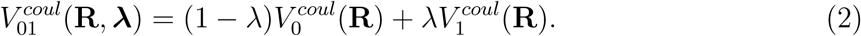

Note that other potentials, such as Lennard-Jones (LJ) potential, whose contributions to proton binding free energy are negligible compared to Coulomb interactions, were not taken into account in this interpolation in the current implementation of GROMACS-cphmd. Beyond this charge interpolation, the total potential energy *V* (**R**, ***λ***) includes three additional *λ*-dependent terms: quantum mechanical correction (*V ^MM^*), double-well biasing potential (*V ^dwbias^*), and pH-dependent term (*V ^pH^*).

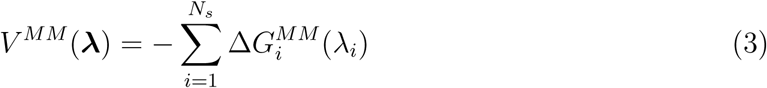

This quantum mechanical correction term compensates for missing quantum effects in classical force fields to make the interpolated potential function flat if the titratable site i is in its reference state. The determination of this potential is carried out by assessing the deprotonation free energy of the single residue in water (reference state) at the force field level. The analytical expressions for the gradients of *V ^MM^* (***λ***) were derived by fitting polynomials to the derivatives of the reference free energy along the charge-interpolation path.

The double-well biasing potential reads:

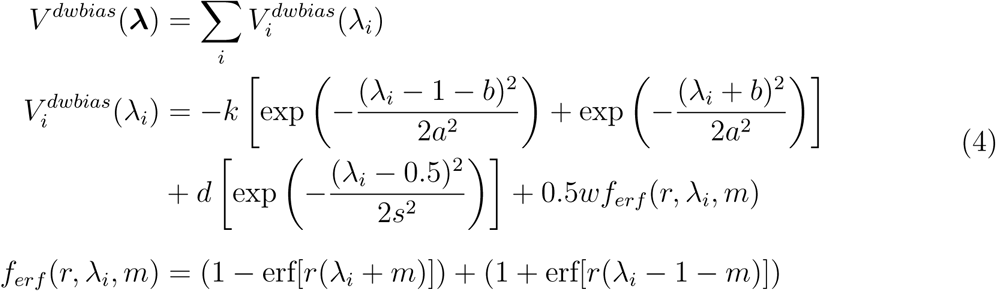

With parameters *k* = 4.7431, *a* = 0.0435, *b* = 0.0027, *d* = 3.75, *s* = 0.30, *w* = 1000.0, *r* = 13.5, *m* = 0.2019, this potential promotes sampling of the *λ* = 0 (protonated) and *λ* = 1 (deprotonated) states by imposing a 7.5 kJ/mol energy barrier between them.^45^ The first two Gaussian terms anchor the potential wells at the end states, the central Gaussian term creates a barrier at *λ_i_* = 0.5, and the error function terms enforce sharp transitions, ensuring efficient exploration of physical protonation states.

The pH-dependent term *V* ^pH^(***λ***) which couples the simulation to the experimental pH by modeling the proton chemical potential, is expressed as a summation over individual contributions from titratable sites. Mathematically, it is given by:

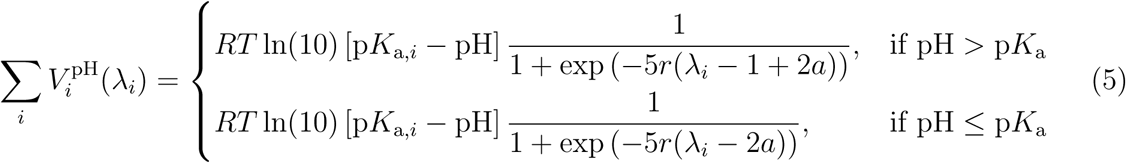

In this expression, p*K*_a*,i*_ represents the experimentally determined p*K*_a_ value for residue i, and the piecewise - defined functions act as a step-like modulation (through the exponential terms) to account for the pH-dependent protonation behavior. Essentially, this term links the simulation to the experimental pH condition by incorporating the influence of proton chemical potential.

Together, these terms collectively enable accurate protonation equilibria by addressing energetic inaccuracies, enhancing sampling efficiency, and incorporating pH effects. Further details are provided in.^35^

### 2.3 System Setup for Standard MD and CpHMD Simulations

Atomistic models for MotAB in the plugged and unplugged states were constructed based on the refined cryo-EM structures of *Campylobacter jejuni* MotAB.

These refined structural models feature enhanced cryoEM density for the N-terminal tail (from C10 to T40) of the MotB dimer which was invisible in previous models (PDB ID: 6YKP and 6YKM, respectively). This improvement allows us to capture previously unresolved atomic details, including the sidechains of residues preceding K15 in MotB, such as glutamic acid at position 14 (E14).

These models were successively inserted into a membrane system within a periodic box (11.6 x 11.6 x 12.6 *nm*^3^) consisting of 80% POPE and 20% POPG, mimicking typical bacterial membrane composition, using the CHARMM-GUI web server.^46,47^ The protein-membrane systems were solvated with TIP3P water and 0.15 M NaCl. Titratable sites were defined, and initial protonation states were assigned using pH-Builder,^48^ which also generated the topology files. A total of 189 titratable residues were determined by designating Glu, Asp, and His residues as titratable (it should be noted that Arg and Lys were not treated as titratable in this research). Buffer particles were added to achieve charge neutrality in the systems. After that, the generated topology files and coordinates were manually inspected and adjusted or processed using in-house scripts. For proteins, the modified CHARMM36m-cph force field was applied,^35,44^ while lipid parameters were borrowed from the original CHARMM36m force field^49,50^ due to the absence of predefined parameters for POPG and POPE in the CHARMM36m-cph force field.

Energy minimization was conducted using the steepest descent algorithm with a tolerance of 1,000 kJ/mol · nm. The LINCS algorithm was employed to constrain covalent bonds involving hydrogen atoms, permitting a time step of 2 fs for molecular dynamics integration.

Electrostatic interactions were calculated using the particle-mesh Ewald (PME) method with a cut-off of 1.2 nm. The Verlet grid cut-off scheme was used for neighbor searching, also with a 1.2 nm cut-off for short-range interactions. Van der Waals interactions were computed with a 1.2 nm cut-off, using a smooth switching function starting at 1.0 nm. Periodic boundary conditions were applied in all spatial dimensions. During equilibration, the temperature was maintained at 300 Kelvin using a v-rescale thermostat with a time constant of 0.5 ps. To accommodate the membrane-protein nature of the system, the pressure coupling type was modified to semi-isotropic, consistent with the default settings provided by pH-Builder. Production simulations were performed across a pH range of 1 to 12, with multiple 100 ns replicas and six parallel groups generated for each pH condition. In the simulations, the titratable residues, including Glu, Asp, and His, were initially assigned to their protonated states.

It is worth noting that although the default two-step NVT–NPT equilibration protocol was successfully employed in the present study and has previously been applied to ligand-gated ion channels,^37^ it is not intrinsically optimized for membrane protein systems. We refined the equilibration workflow by implementing a six-step procedure, analogous to the membrane protein equilibration protocol provided by CHARMM-GUI, and observed a modest improvement in convergence performance with this optimized protocol (Fig. S1). The corresponding molecular dynamics parameter files are provided on GitHub (see link in the “Code Availability” subsection).

Convergence is checked by monitoring the average fraction of proton dissociation (*f_D_*) as a function of time by including all frames up to the current time point. Here, *λ* values are dynamically updated during the simulation according to the local physical interactions. *f_D_*for a residue is subsequently calculated based on the *λ* values. Specifically, a residue is considered protonated when *λ* ≤ 0.2 and deprotonated when *λ* ≥ 0.8. *λ* values between 0.2 and 0.8 are excluded as non-physical samples. For residues with a single titratable site (such as Asp and Glu), *f_D_* is calculated as:

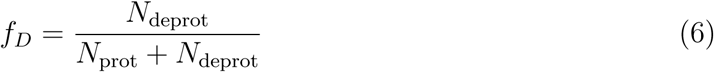

where *N*_prot_ and *N*_deprot_ are the counts of frames in the protonated and deprotonated states, respectively. For His with two titratable sites, its fraction of proton dissociation *f ^His^* is calculated as:

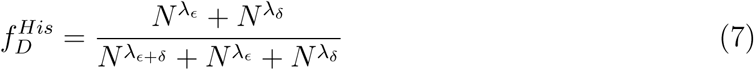

where *ɛ* and *δ* refer to single protonation at *N_ɛ_*and *N_δ_* atoms of the His residue, respectively, and *ɛ* + *δ* refers to protonations on both sites. *N^λɛ^*^+^*^δ^*, *N^λɛ^*, and *N^λδ^* are the numbers of frames in which *λ_ɛ_*_+*δ*_ *>* 0.8, *λ_ɛ_ >* 0.8, and *λ_δ_ >* 0.8, respectively. The p*K*_a_ value for each residue is subsequently calculated using a variant of Henderson–Hasselbalch equation:

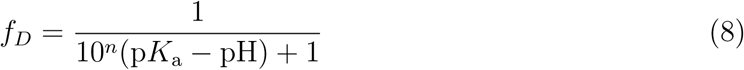

where *n* is the Hill coefficient, which quantifies the degree of coupling between residues. The error of *f_D_* is calculated by the method of block analysis. For each trajectory, we split it into 2, 4, 8, 16, 64, 128, and 256 blocks, then calculate the standard error, employing the block averaging method to locate a plateau region in which the error remains stable.^51^ In cases where such a plateau cannot be identified, the result obtained from the maximum block size should be used.

To investigate how the rotation of MotAB is governed by the presence of the plug region and the protonation states of D22, four distinct models were constructed on the basis of the refined cryo-EM structures of CjMotAB in its plugged and unplugged conformational states. (1) The first model was a plug-removed variant of MotB with residues N41 to F56 truncated, derived from the refined plugged-state MotAB structure; in this model, F-D22 was protonated, whereas G-D22 remained deprotonated. (2) The second model was an identical plug-removed variant, with D22 residues in both the F and G chains retaining their deprotonated state. (3) The third model was also based on the plug-removed variant, with F-D22 being deprotonated and G-D22 being protonated. (4) The last model corresponded to the unplugged conformational state, in which both D22 residues were maintained in a protonated form. All four models were built using CHARMM-GUI with parameters consistent with those utilized in the accompanying CpHMD simulations. Subsequent MD simulations were executed in GROMACS with the same equilibrium protocol. The simulation temperature was stabilized at 300 K via a v-rescale thermostat with a time constant of 1.0 ps, and each independent simulation was run for a total duration of 1000 ns (Table S2).

### 2.4 Comparison with Other p*K*_a_ Predictors

The p*K*_a_ values obtained from CpHMD simulations were evaluated against predictions from three established single-structure methods: DeepKa,^40–42,52^ PropKa 3.0^38^ and PypKa.^39^ DeepKa represents a deep-learning approach to p*K*_a_ prediction, while PropKa 3.0 employs an empirical framework that incorporates desolvation effects and dielectric response in its calculations. And PypKa calculates p*K*_a_’s by solving Possion-Boltzmann equation numerically.

To further cross-validate the accuracy of the p*K*_a_ values for key Asp residues, Free Energy Perturbation (FEP) calculations were employed.^53–55^ A perturbation was applied to the side chain of Asp residues (Fig. 2E). Based on FEP, the free energy difference Δ*G* associated with the deprotonation process can be computed.

**Figure 2:**
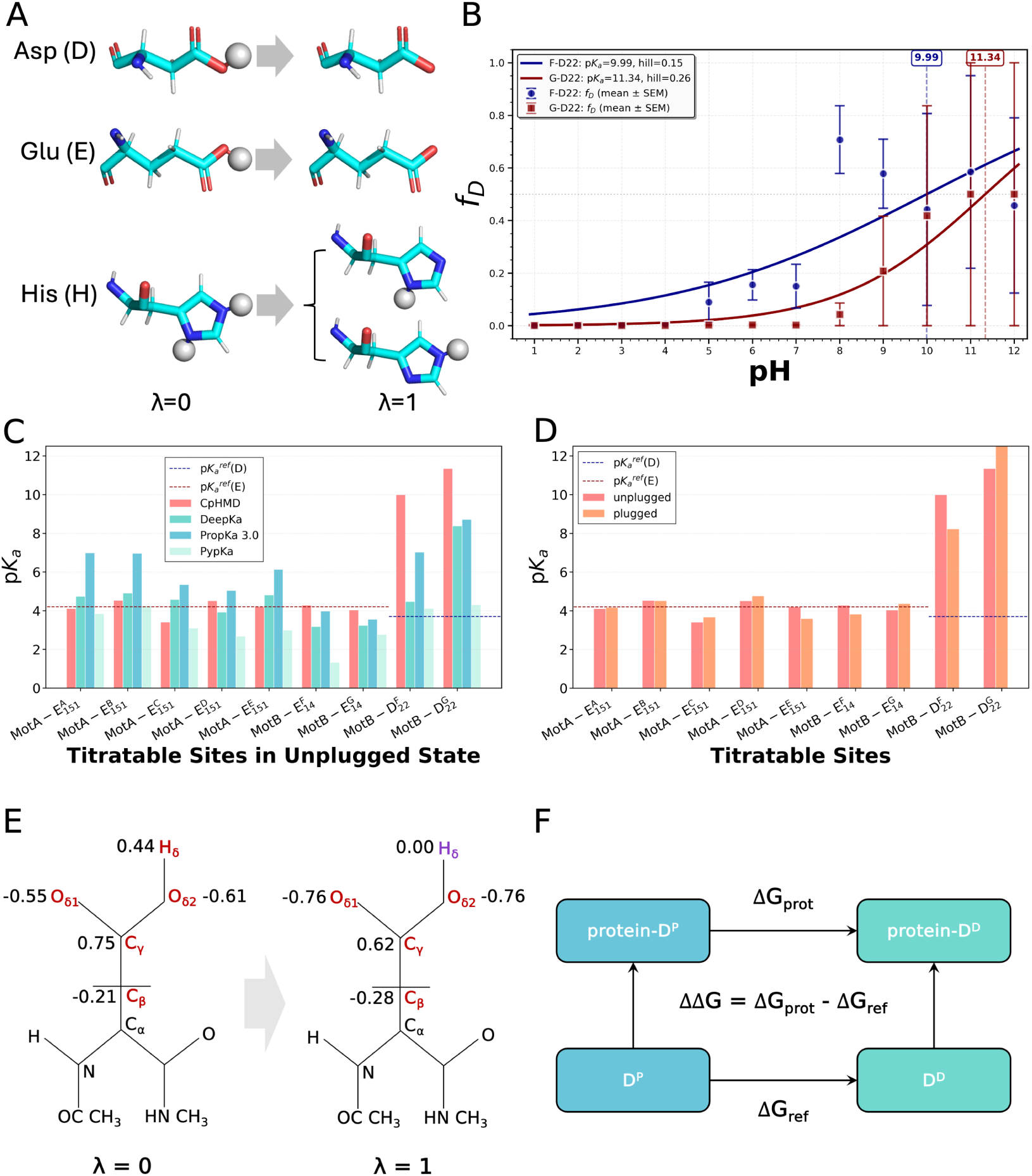
**Protonation thermodynamics and pK modulation of titratable residues in CjMotAB**. (A) Schematic representation of the alchemical transformation (*λ* = 0 →*λ* = 1) for aspartate (Asp, D), glutamate (Glu, E), and histidine (His, H) residues, used to compute the free energy of protonation in the protein environment relative to bulk solution. (B) Titration curves of MotB F- and G-D22. F chain: Hill fit = 0.15, p*K*_a_ = 10.0. G chain: Hill fit = 0.26, p*K*_a_ = 11.3. *f_D_* = fraction of deprotonated residue. (C) Comparison of p*K*_a_ values for titratable sites in the CjMotAB system, calculated using CpHMD, DeepKa, PropKa 3.0, and PypKa. Reference p*K*_a_ values are standard for Glu (4.2) and Asp (3.7) in aqueous solution.^67^ (D) CpHMD-derived p*K*_a_ values and shifts under plugged and unplugged conformations. (E) Partial charge distributions of an Asp residue in its protonated (*λ* = 0) and deprotonated (*λ* = 1) states, as defined by the CHARMM36m-cph force field for FEP calculations. (F) Thermodynamics cycle for protein-specific p*K*_a_ calculations.

FEP calculations were performed using GROMACS with the following parameters: The free energy module was enabled, and simulations were initiated from the initial *λ* state (*λ* = 0). Smooth transitions between protonated and deprotonated states were achieved using either 11 or 21 evenly spaced *λ* values, ranging from 0.00 to 1.00 in 0.05 or 0.10 increments to ensure efficient overlap evaluations. Energy minimization was first performed, followed by six sequential equilibration steps. These stages involved a gradual reduction of restraining forces on various system components and a progressive increase in surface tension along the membrane xy-plane. Temperature was maintained at 300 K using a V-rescale thermostat with a time constant of 1.0 ps. Additionally, other basic parameters were consistent with those of the CpHMD simulations. Soft-core potentials were employed to avoid singularities during the transformation, with a soft-core alpha parameter of 0.3 and a soft-core sigma value of 0.25. Coulomb interactions were linearly interpolated during the transformation. Finally, the Multistate Bennett Acceptance Ratio (MBAR) was utilized to compute the free energy differences.^56,57^

A thermodynamic cycle was constructed to quantify shifts in p*K*_a_ values,^58^ with Asp residues in aqueous solution serving as the reference system. The calculation of the equilibrium constant *K_a_* for each reaction essentially entails the determination of the standard reaction free energy Δ*G*:

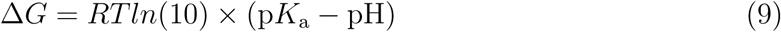

From this, we derive:

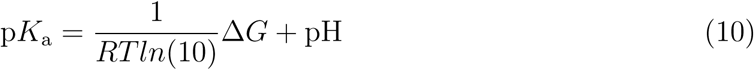

and

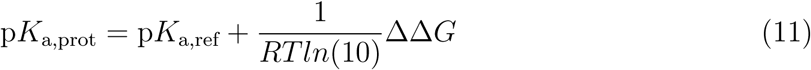

where p*K*_a,prot_ and p*K*_a,ref_ denote the p*K*_a_ values of the titratable group in the protein and the reference molecule (Asp) in aqueous solution, respectively; ΔΔ*G* = Δ*G*_prot_ − Δ*G*_ref_ represents the double free energy difference corresponding to the thermodynamic cycle.

### 2.5 Conformational Dynamics Analysis and Hydrogen Bond Networks

To characterize the rotational dynamics of MotA relative to MotB, a path collective variable (CV) was defined.^59,60^ The progress variable *s* of this path CV is a one-dimensional order parameter, which quantifies both the system’s progression along a predefined reference pathway and its deviation from this pathway, and is mathematically expressed as:

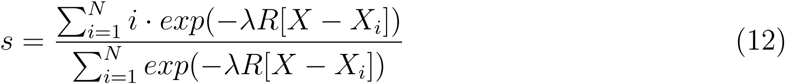

Here, *X* represents the coordinates of the instantaneous protein conformer sampled during simulations. The similarity between *X* and each reference frame *X_i_* is quantified using a distance metric *R*[*X* − *X_i_*] (specifically, RMSD). *λ* serves as a smoothing parameter and is set to be 103.4 and 175.1 for the unplugged model and the plugged model, respectively. The progress variable *s* ranges from 1 to N (with N=72 in this study), spanning the full rotational range of MotA from the initial state (0°) to the final state (72°). This final rotational state was computationally achieved by reassigning chain identifiers. The reference pathway for the path CV was constructed via linear morphing in PyMOL, where each consecutive frame corresponds to a 1° incremental rotation of MotA. This discretized pathway enables the PLUMED driver to map the rotational state of MotA throughout the simulations.

To rationalize the observed p*K*_a_ shifts of MotB D22 in the unplugged conformational state, MDAnalysis^61,62^ was additionally utilized to quantify three environmental descriptors of this residue: (1) the spatial distance between MotB D22 and clusters of highly hydrophobic residues; (2) the solvent-accessible surface area (SASA); and (3) the number of water molecules in its immediate vicinity.

Comprehensive interaction analysis between different residues in the MotAB complex was performed using GetContacts (https://github.com/getcontacts/getcontacts). This analysis included the identification of interactions between MotB D22 and MotA residues (L158, T189), characterization of the interaction between MotA E151 and MotB E14/K15, and examination of less frequently reported interactions, including those between MotA A161 and MotB D22, and between MotA Y197 and MotA E151. The structural basis for these interactions was analyzed to understand their potential functional implications.

To investigate the proton transfer pathway, the hydrogen bond network originating from the titratable oxygen atom of MotB F-D22 was analyzed. MDAnalysis was employed for trajectory analysis, and a breadth-first search (BFS) algorithm^63^ was implemented to systematically identify all possible hydrogen bond pathways up to five grades. And a hydrogen bond is defined such that the distance between the acceptor and the donor does not exceed 3.5 Å, and the acceptor–hydrogen–donor bond angle is no less than 120 degrees.

The hydrogen bond network was defined with specific criteria: first-grade hydrogen bonds were direct donor-acceptor links between MotB D22 and neighboring residues or water molecules; second-grade hydrogen bonds were subsequent donor-acceptor connections formed by the first acceptor acting as a new donor; and higher-grade hydrogen bonds continued this network identification process. Pathways terminating in water molecules were filtered out to focus on biologically relevant interactions. The stability of the hydrogen bond network was assessed throughout the simulation trajectories to evaluate its potential role in proton transfer via the Grotthuss mechanism.

### 2.6 Sequence Conservation and Coevolution Analysis with EV-couplings

The conservation of these key residues identified in CpHMD and their associated features was evaluated using multiple sequence alignment (MSA) and coevolution analysis with EV-couplings.^64–66^ Rooted in the Potts statistical mechanics model, EVcouplings quantifies the impact of amino acid mutations through a defined energy function. This function consists of two core components: site-specific bias parameters (*h_i_*) that describe the intrinsic propensity of individual residues, and pairwise coupling parameters (*J_ij_*) that capture the coevolutionary interdependence between residue pairs i and j. The total energy of a given sequence configuration *σ* is formulated as:

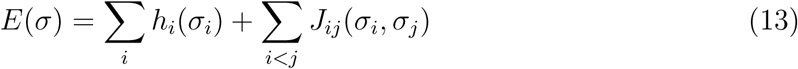

Two distinct model variants were employed for comparative analysis: an independent model that incorporates only the site-wise bias term (*h_i_*), and an epistatic model that accounts for both the *h_i_* and *J_ij_*terms to characterize residue co-dependencies.

### 2.7 Code Availability

The source code utilized in this work and the custom scripts developed for subsequent data analysis are publicly available in the following GitHub repository: https://github.com/luojc712/MotAB-rotation-mechanism.

## 3 Results

### 3.1 Refined Model of CjMotAB in Unplugged State using Cryo-EM

As previously reported, single-particle cryo-EM was used to resolve two CjMotAB variants: a plug-containing form, in which the MotB plug helix mediates self-inhibition, and an unplugged variant lacking the plug helix to mimic the activated state, at resolutions of 3.1 Å and 3.0 Å, respectively.^10^ Building on these datasets, we leveraged improvements in the latest version of CryoSPARC to further enhance structural resolution. Particles were repicked from denoised micrographs, and the conventional two-dimensional (2D) classification step was omitted. Instead, particles were directly subjected to three-dimensional (3D) classification to eliminate noise and poorly resolved classes.^43^ The selected particles were subsequently refined using reference-based motion correction to achieve higher-resolution reconstructions. This optimized workflow improved the cryo-EM map resolutions to 2.6 Å for the plugged state and 2.3 Å for the unplugged state (Table S1). The resulting structural models exhibited substantial improvements in refinement quality, including increased model completeness, more accurate side-chain conformations, and clearly resolved bound water molecules (Fig. S2). Notably, previously ambiguous electron density in the MotB N-terminal tail (residues C10–T40) was resolved, allowing detailed modeling of side chains, including residues such as E14 (adjacent to K15) that were not characterized in earlier studies.^31^

### 3.2 CpHMD-Derived p*K*_a_ Predictions for CjMotAB Key Residues and Cross-Validation

Using these refined high-resolution structures, we performed CpHMD simulations to examine how protonation state changes of key residues are coupled to conformational dynamics.

We first focused on the active, unplugged state of CjMotAB. Given the established role of D22 in driving stator unit rotation, our analysis centered on its titration behavior (titration profiles for additional residues are shown in Fig. S3, with convergence analyses provided in Figs. S4–S5). The results revealed significant p*K*_a_ upshifts for D22 in the MotB dimer relative to reference p*K*_a_ values,^67^ with estimated values of 10.0 and 11.3 for F-D22 (MotB D22 in the F chain) and G-D22 (MotB D22 in the G chain), respectively. These elevated values indicate that both D22 residues remain protonated under physiological conditions.

To validate these findings, we compared CpHMD-derived p*K*_a_ values with predictions from DeepKa, PropKa 3.0, and PypKa (Fig. 2C). For MotA residue E151—previously proposed to play a critical role in rotation—PropKa 3.0 predicted a substantial p*K*_a_ upshift, whereas CpHMD and DeepKa did not support this trend. In addition, all methods consistently indicated only minor p*K*_a_ perturbations for MotB E14, suggesting that this residue remains stably deprotonated. Correlations among the three prediction methods under both conformational states are presented in Figs. S6–S8. In contrast, PypKa did not predict significant p*K*_a_ shifts for any of the residues of interest.

To investigate how conformational changes affect protonation equilibria in MotAB, we additionally performed CpHMD simulations over a pH range of 1–12 for the inactive, plugged conformation, using the refined cryo-EM structure as the starting model. The simulations yielded a p*K*_a_ of 8.2 for F-D22 and an exceptionally high p*K*_a_ for G-D22 (exceeding 12). Both residues exhibit pronounced upward shifts, consistent with the trend observed in the unplugged state.

### 3.3 Free Energy Perturbation Calculations to Cross-Validate MotB D22 p*K*_a_ Shifts

Our CpHMD simulations indicate that both D22 residues exhibit notably elevated p*K*_a_ values, as expected—this finding supports the previous hypotheses. In contrast, other fast but potentially less accurate predictors (such as PypKa) failed to detect these p*K*_a_ shifts. To further validate the CpHMD-derived p*K*_a_ values, we performed free energy calculations using a thermodynamic cycle (Fig. 2F) and free energy perturbation (FEP) to quantify the double free energy differences (ΔΔ*G*) associated with deprotonation.

For F-D22, the calculated ΔΔ*G* of deprotonation was 12.69 kcal/mol, corresponding to a p*K*_a_ shift of 9.2. In contrast, G-D22 exhibited a much larger ΔΔ*G* of 36.98 kcal/mol, corresponding to an extraordinary p*K*_a_ shift of 26.93. This marked difference between the two homologous residues reveals a pronounced functional asymmetry within the complex. The significantly higher deprotonation barrier for G-D22 suggests that it is far less likely to release a proton under physiological conditions compared to F-D22, with potential implications for directional proton transfer and the overall rotation cycle of the motor.

Overall, while such high ΔΔ*G* values may arise from the challenge of accurately modeling the dielectric constant in the membrane protein environment,^68,69^ these results consistently indicate that both D22 residues have a substantially higher free energy barrier for proton release than typical Asp residues in aqueous environments, and the barrier in the G chain is significantly higher than that in the F chain (Table S3–S5, Fig. S9, S10).

### 3.4 D22 Protonation Regulates MotA Rotational Dynamics

All p*K*_a_ prediction methods consistently revealed pronounced upshifts for D22 in both the unplugged and plugged conformations, underscoring its critical role in MotAB rotation. To further elucidate the underlying mechanism, we analyzed the rotational dynamics of MotAB, focusing on the relationship between D22 protonation states and conformational motion.

Analysis of CpHMD trajectories at pH 7.0 revealed a clear contrast between the two conformations (Fig. 3A–B). The unplugged state displayed substantial rotational dynamics, with a maximum rotation angle of approximately 20° and an average of 8.7°. In contrast, the plugged state exhibited only minimal angular displacement, consistent with its self-inhibited nature.

**Figure 3:**
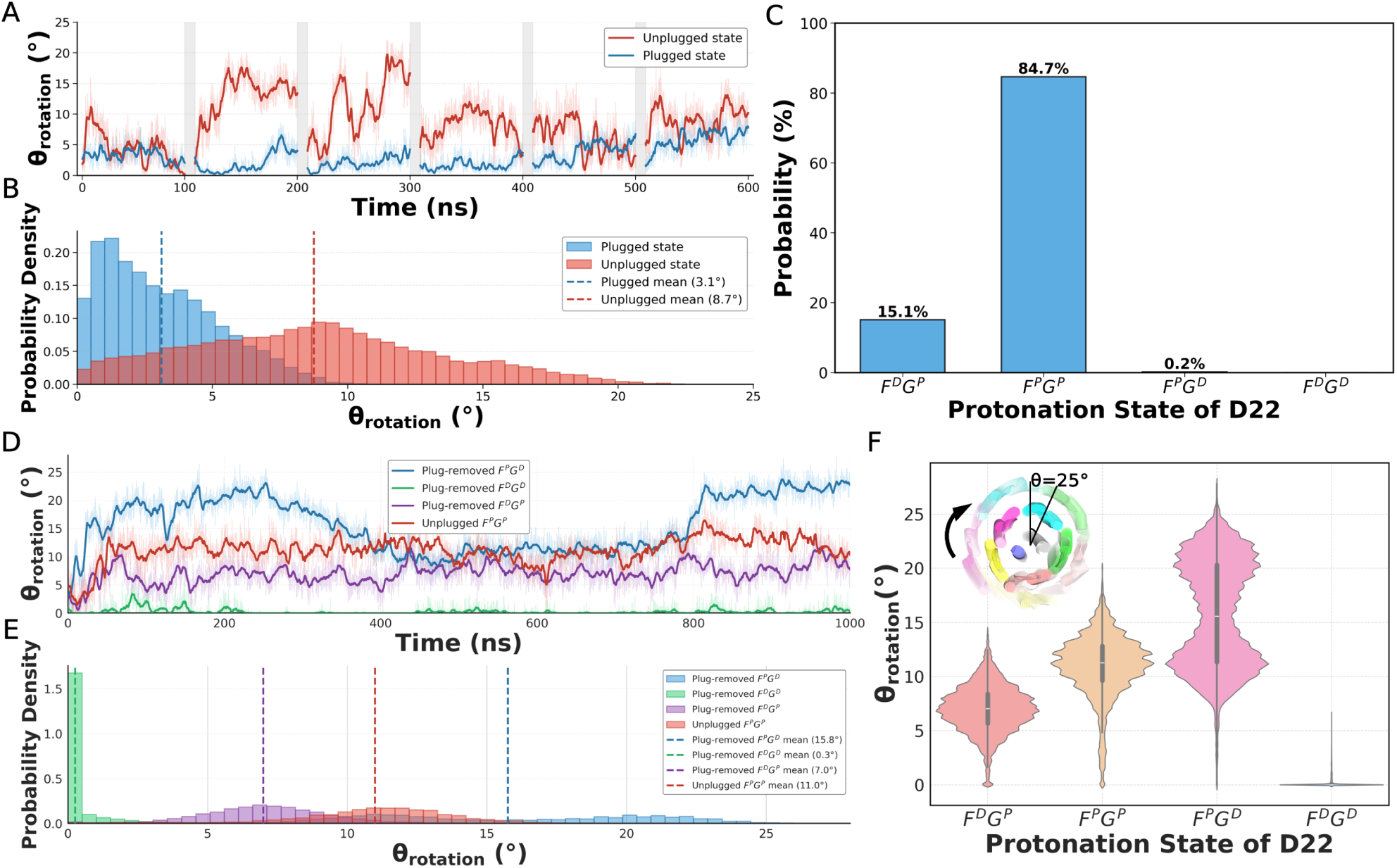
Rotational dynamics of MotA and its correlation with the protonation states of D22. (A) Time evolution of the rotation angle for the plugged and unplugged MotAB conformations. The plot combines six independent 100 ns replicate CpHMD simulations, with gray gaps indicating the boundaries between individual replicates. (B) The corresponding distributions of MotA rotation in the plugged and unplugged states. (C) The probability of sampling each D22 protonation state combination of the F and G chains in CpHMD simulations. Here, the superscript “P” indicates a protonated state and “D” indicates a deprotonated state; the first character corresponds to the F-chain D22 residue, and the second to the G-chain D22 residue. (D, E) Rotation angles of the plugged (removing plug) and unplugged MotAB conformations over time in standard MD simulation. (F) Angle distribution of each protonation state combination of D22 of the F and G chains in standard MD simulations.

We next evaluated the population of different protonation state combinations sampled during the CpHMD simulations. The *F^P^ G^P^*, *F^D^G^P^* and *F^P^ G^D^* states—where superscript “P” denotes a protonated state and “D” denotes a deprotonated state, with the first and second letters referring to the F and G chain D22 residues, respectively—were found to dominate, with relative populations of 84.7%, 15.1%, and 0.2% (Fig. 3C). Notably, the fully deprotonated *F^D^G^D^* state was not observed, suggesting that it is unlikely to represent a stable or biologically relevant configuration. Moreover, all sampled states except *F^D^G^D^* exhibited rotational motion (Fig. S11).

### 3.5 Plug Removal and D22 Protonation: Dual Requirements for MotAB Rotation

While prior structural and functional studies have established that the plugged conformation inhibits rotation and that unplugging is necessary for MotAB function, which aligns well with our simulation results, these experiments cannot resolve the protonation-dependent driving forces underlying rotation. Our CpHMD simulations extend these findings by revealing a strong coupling between G-D22 protonation and MotA rotation, suggesting that protonation switching constitutes a critical regulatory element beyond plug removal. This raises a key question: is unplugging alone sufficient to initiate rotation, or is D22 protonation also required?

To address this, we constructed models derived from the cryo-EM plugged structure with the plug region removed and performed 1000 ns standard MD simulations under fixed protonation conditions. Three protonation states were examined: the plug-removed *F^P^ G^D^* model (F-D22 protonated, G-D22 deprotonated), the plug-removed *F^D^G^P^*model (F-D22 deprotonated, G-D22 protonated), and the plug-removed *F^D^G^D^*model (both residues deprotonated). For comparison, we also simulated the cryo-EM unplugged structure with both D22 residues protonated (the unplugged *F^P^ G^P^* model).

Note that although both plug-removed and unplugged models lack the plug region, they originate from different structural templates. The plug-removed models were generated by deleting the plug from the plugged cryo-EM structure, whereas the unplugged model is based directly on the experimentally determined unplugged structure.

As shown in Fig. 3D–F, pronounced rotational motion was observed in both the plug-removed *F^P^ G^D^* and *F^D^G^P^* models, comparable to that seen in the unplugged *F^P^ G^P^* model, which represents the expected active state. In contrast, the plug-removed *F^D^G^D^* model exhibited no detectable rotation, indicating that removal of the plug alone is insufficient to enable MotAB rotation when both D22 residues are deprotonated.

Together, these results demonstrate that CjMotAB rotation requires two essential conditions: displacement of the plug and protonation of at least one MotB D22 residue.

### 3.6 Asymmetric Hydration of D22 and Water Accessibility Regulation of D22 Protonation

Although the differences in p*K*_a_ between the two D22 residues may partly arise from their distinct local interactions with neighboring MotA residues (Fig. S12), which differentially stabilize the protonated state, we further investigated the role of solvation in shaping their protonation behavior. Given the importance of water molecules in proton transfer, we examined the hydration environment surrounding MotB D22. As shown in Fig. 4A, water molecules are distributed asymmetrically around the D22 sites in the F and G chains.

**Figure 4:**
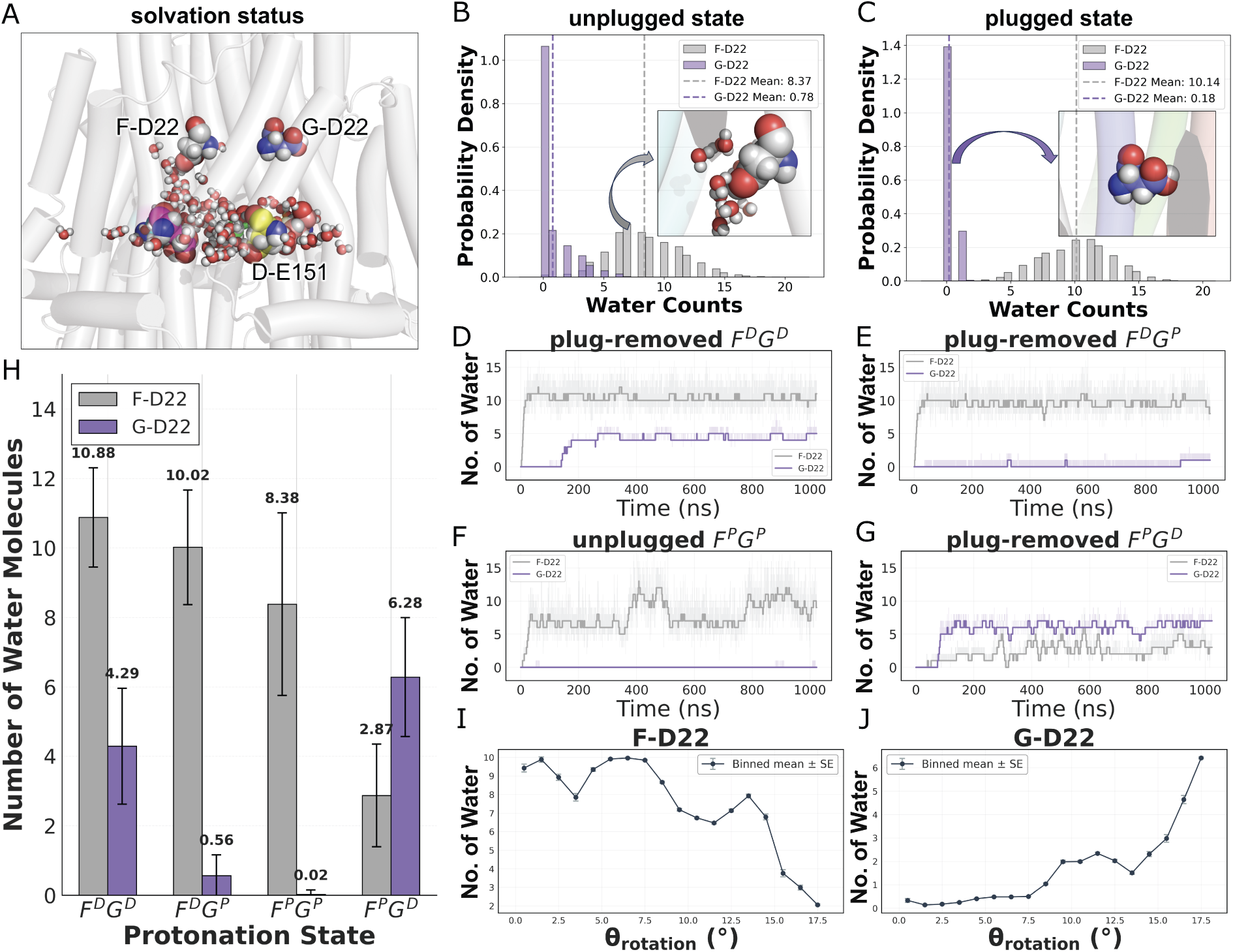
Hydration and hydrogen bonding network analysis of MotB D22 in the active state of MotAB. (A) 3D structural schematic diagram of the asymmetric water molecule distribution on both F- and G-D22. (B, C) Distribution of water molecular counts around MotB D22 side chain within 5 Å in both plugged and unplugged state CpHMD simulations. (D, E, F, G) Water molecular counts around F- and G-D22 in standard MD simulations over time. *F^D^G^D^* corresponds to *F^D^G^D^* in the plug-removed model. *F^D^G^P^* corresponds to *F^D^G^P^*in the plug-removed model. *F^P^ G^P^* corresponds to *F^P^ G^P^*in the unplugged model. *F^P^ G^D^* corresponds to *F^P^ G^D^* in the plug-removed model. (H) Statics of water molecular counts around MotB D22 in standard MD simulations. (I, J) Changes in hydration of F- and G-D22 during the rotation process.

To quantify hydration, we monitored the number of water molecules within 5 Å of each D22 residue during CpHMD simulations of both unplugged and plugged conformations at pH 7.0. A striking asymmetry was observed: F-D22 remains highly hydrated in both states, with average coordination numbers of approximately eight and ten water molecules, respectively. In contrast, G-D22 is largely dehydrated, with average counts of only one and zero water molecules in the unplugged and plugged states, respectively (Fig. 4B–C). These results reveal a clear correlation between hydration and p*K*_a_ shifts: increased dehydration of the D22 microenvironment corresponds to higher p*K*_a_ values and larger barriers to deprotonation.

To further probe the coupling between hydration and protonation, we analyzed standard MD simulations by tracking water occupancy of D22 during transitions between deprotonated and rotationally dominant protonated states. As shown in Fig. 4D–G, simulations initialized without nearby water molecules demonstrated rapid solvent infiltration into the MotB transmembrane region during equilibration. The resulting hydration level depends strongly on protonation state: the deprotonated form of D22 consistently exhibits a more extensive hydration shell than the protonated form. Analysis of average water coordination further highlights distinct chain-specific behaviors (Fig. 4H).

For F-D22, hydration progressively decreases as the residue transitions from the deprotonated state to the protonated state, and declines further upon adopting the protonated conformation associated with rotation. In contrast, G-D22 displays a cyclic hydration pattern: water occupancy decreases upon protonation and during the rotation-associated state, but increases again upon returning to the deprotonated state.

We further examined how rotation-induced conformational changes influence hydration at the D22 sites. As shown in Fig. 4I–J and Fig. S13, MotA rotation reduces hydration around F-D22 while increasing hydration around G-D22, indicating that conformational dynamics directly reshape the local solvent environment of these residues.

Collectively, these findings demonstrate that D22 protonation is tightly coupled to its local structural context and hydration dynamics, highlighting a coordinated interplay between solvent accessibility, protonation state, and conformational transitions throughout the rotational cycle.

### 3.7 D22 Sidechain Rotation and Protonation Coupling

Previous studies based on cryo-EM structures (including the refined structure from this work) have noted that the side-chain conformations of MotB D22 exhibit significant dynamics across different structural states.^10^ To further investigate this, we analyzed the distribution of the *χ*_1_ dihedral angle (N–C*_α_*–C*_β_*–C*_γ_*) of the MotB D22 side chain during standard MD simulations.

Our results reveal that in simulations exhibiting minimal rotational motion, the *χ*_1_ dihedral angle of the F-chain D22 side chain primarily adopts a *gauche* conformation (cos(*χ*_1_) ≈ 0.5), while G-chain D22 remains stable in a *trans* conformation (cos(*χ*_1_) ≈ −1.0). In contrast, in simulations characterized by significant rotational movement, the D22 residues in either the F or G chains undergo dynamic transitions between the *gauche* and *trans* states (Fig. 5A–D).

**Figure 5:**
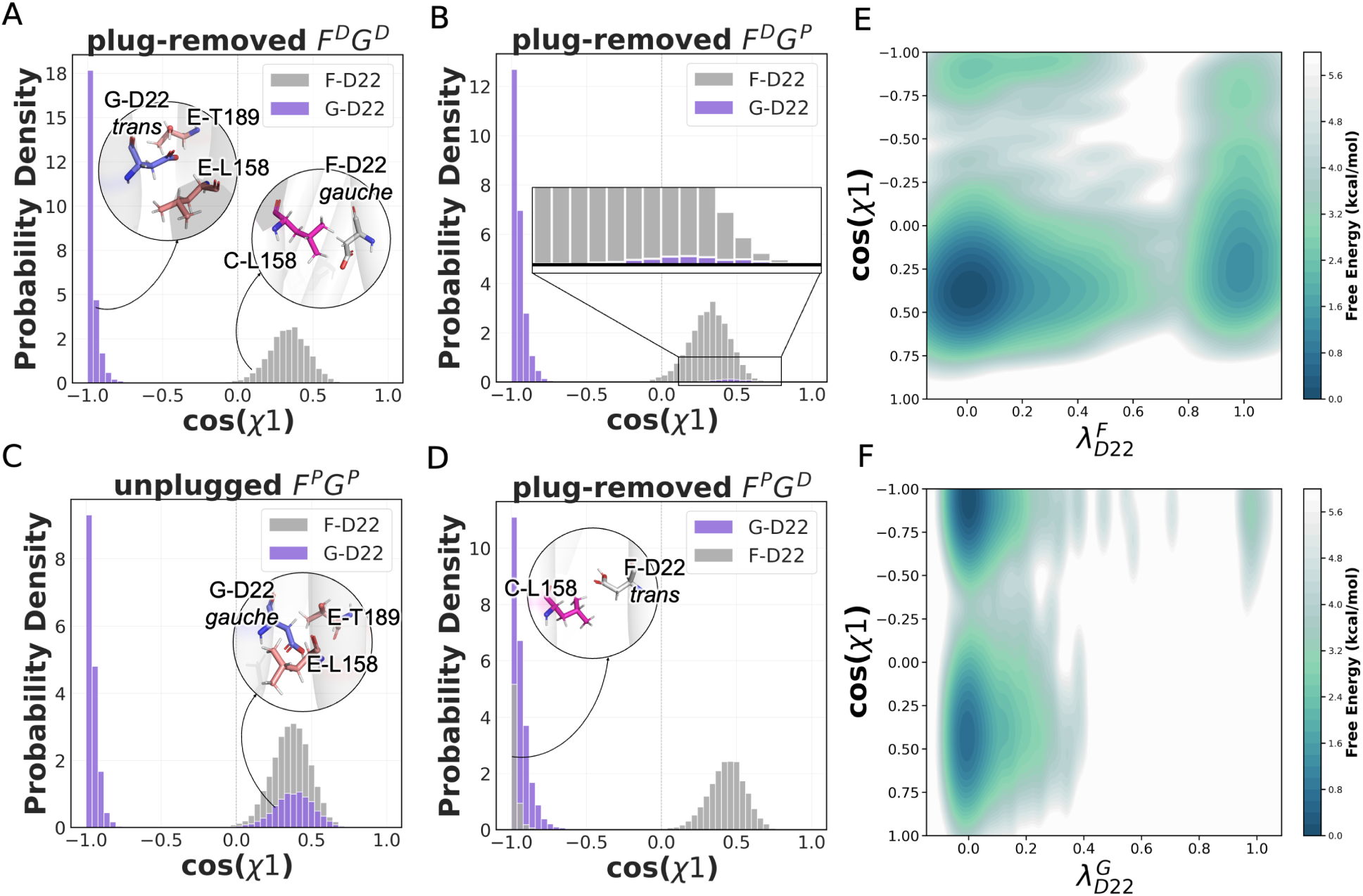
Side chain dihedral angle dynamics and rotation mechanism. (A-D) Distributions of the D22 side-chain *χ*_1_ dihedral angle from standard MD simulations of the plug-removed and unplugged states under distinct D22 protonation conditions: (A) *F^D^G^D^*, (B)*F^D^G^P^*, (C)*F^P^ G^P^*, and (D) *F^P^ G^D^*. (E, F) Free energy surfaces as a function of the D22 side chain dihedral angle *χ*_1_ and its protonation states in unplugged state CpHMD simulations under pH 7.0.

To establish the direct relationship between protonation state and *χ*_1_ dihedral dynamics, we analyzed the results of the CpHMD simulations of the unplugged state at pH 7.0 and constructed two-dimensional free energy surfaces as a function of the protonation state (*λ*) and cos(*χ*_1_) (Fig. 5E–F).

For the G-D22, two energetically favorable basins emerged at *λ* ≈ 0 (protonated), corresponding to both *gauche* and *trans* conformations. By contrast, at *λ* ≈ 1 (deprotonated), only a single, narrow low-energy region was present, exclusively favoring the *trans* conformation. These results are fully consistent with our standard MD observations.

A similar trend was observed for the F-D22. Notably, the low-energy trans conformation of the F-D22 sampled at *λ* ≈ 1 likely corresponds to the subsequent half-cycle, during which the functional roles of the F- and G-D22 residues are exchanged.

Overall, these findings demonstrate that in models capable of rotation, at least one protonated D22 residue exhibits dynamic conformational shifts in its side-chain *χ*_1_ dihedral angle. Conversely, in non-rotating models, these two residues remain stable in their respective low-energy conformations.

### 3.8 Residue Interaction Analysis Reveals Asymmetric Interactions Mode of D22

Further contact analysis revealed the relationship between the interactions of MotB D22 with surrounding residues and its conformational changes. In CpHMD simulations of the unplugged state conformation, we observed a side-chain hydrogen bond interaction between G-D22 and E-T189, whereas no such stable hydrogen bond existed for F-D22 (Fig. 6A–B, Fig. S14). This is highly likely responsible for the conformational differences between the two MotB D22 sites. We further analyzed the trajectories from four sets of standard MD simulations to examine the relationship between the conformation of MotB D22 and the hydrogen bonds formed with MotA T189.

**Figure 6:**
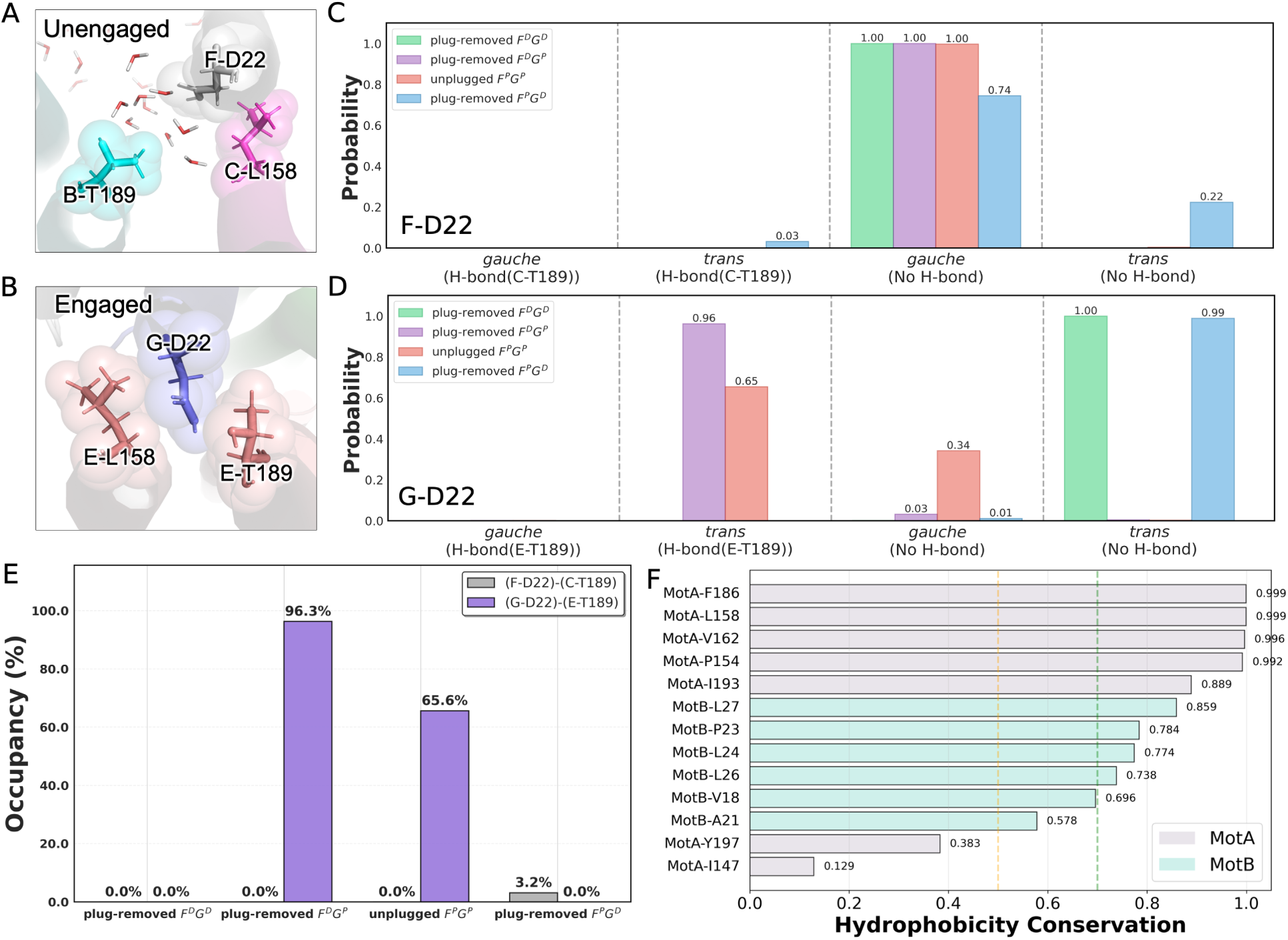
Analysis of residue interactions in the MotAB stator unit. (A, B) Three-dimensional structural representations of key residues interacting with MotB D22; residues are shown in sphere and stick format, labeled with identifiers. (C, D) Relationship between the *χ*_1_ side-chain dihedral angle conformation of MotB D22 and hydrogen bonding with MotA T189 in four sets of standard molecular dynamics simulations. (C) represents F-D22 and (D) represents G-D22. (E) Hydrogen bond occupancy between MotB D22 and MotA T189. (F) Hydrophobicity conservation of key residues in the MotAB.

As shown in Fig. 6C–D, MotB D22 consistently adopted the *trans* conformation when engaged in a hydrogen bond with MotA T189. For F-D22, the lack of a hydrogen bond with MotA T189 coincided with the *gauche* conformation. In the case of G-D22, while deprotonated and rotationally static, it retained the *trans* conformation without hydrogen bonding with MotA T189; conversely, upon protonation and rotational displacement, G-D22 either formed a hydrogen bond with MotA T189 or underwent hydrogen bond dissociation accompanied by a conformational shift to *gauche* conformation.

To quantify hydrogen bond stability, hydrogen bond occupancies were calculated. As shown in Fig. 6E, G-D22 formed an extremely stable hydrogen bond with E-T189 in the plug-removed *F^D^G^P^* model, whereas hydrogen bond stability decreased in the unplugged *F^P^ G^P^* model. In the plug-removed *F^P^ G^D^* model, F-D22 also formed transient hydrogen bonds with C-T189.

These results indicate that the formation of hydrogen bonds correlates with conformational changes in the D22 side-chain dihedral angle. For F-D22, hydrogen bond formation induces a *gauche* to *trans* transition in its side-chain dihedral angle. In contrast, for G-D22, hydrogen bond breakage triggers a *trans* to *gauche* transition.

These contact analysis results align with prior mutational experiments.^10^ In *Salmonella enterica*, substitutions at MotA T209 and MotB D33 were shown to impair MotAB-driven flagellar rotational function, as evidenced by a reduced migration radius. Notably, these two residues correspond to MotA T189 and MotB D22 in CjMotAB. Additionally, several important amino acid residues were identified from interaction analysis (Fig. S15-S16), coevolution analysis (Fig. S17-S19), and previous mutagenesis experiments. Fig. 6F shows the hydrophobic conservation of these residues, and we found that most of them exhibit clear hydrophobic conservation. This provides a hydrophobic framework for the interactions between protonated D22 and these residues.

## 4 Discussion

### 4.1 Comparison of p*K*_a_ Predictions

In this study, we combined static structure-based p*K*_a_ prediction tools with molecular dynamics-based CpHMD simulations to evaluate the protonation behavior of titratable residues, enabling a direct comparison across different methodological frameworks. Notably, the PropKa 3.0 prediction for MotA E151 is consistent with previous reports for this system. However, static structure-based approaches—including PropKa—do not fully agree with the CpHMD results, underscoring the limitations of methods that rely on a single conformation. Earlier structural models lacked the N-terminal tail of MotB, leading to the omission of key residues such as E14, which resides in the same charged layer as MotA E151. The absence of this region likely perturbs the local electrostatic environment and introduces systematic errors in p*K*_a_ estimation.

From a methodological perspective, CpHMD offers clear advantages over static predictors such as DeepKa, PropKa 3.0, and PypKa. While these tools are computationally efficient, they cannot capture the dynamic coupling between protonation states and conformational changes, nor do they adequately account for the unique dielectric properties of membrane environments. In contrast, CpHMD explicitly samples protonation states alongside conformational dynamics and solvent interactions, enabling a more realistic description of electrostatics and protonation equilibria.

Nevertheless, CpHMD has its own limitations. For example, van der Waals interactions of protons are retained even in the deprotonated state, which is not strictly physical. However, control calculations indicate that this artifact contributes only 0.2–1.5 kcal/mol to the free energy, and is therefore negligible (Table S3–S5, Fig. S9–S10).

By comparison, FEP methods can more rigorously treat deprotonation by alchemically transforming a proton into a non-interacting dummy particle. However, FEP is typically limited to evaluating one residue at a time and cannot easily capture cooperative protonation behavior across multiple sites. Therefore, for complex membrane proteins such as CjMotAB, CpHMD provides a practical balance between accuracy and efficiency, offering both reliable p*K*_a_ estimates and dynamic mechanistic insight. Future integration with enhanced sampling or coarse-grained approaches^32,70,71^ may further improve the characterization of conformationally coupled protonation dynamics.

### 4.2 Coupling Between Asymmetric Solvation and Protonation of D22

A key finding of this study is the cyclic fluctuation in hydration around D22, which closely tracks its protonation state during the rotational cycle. This asymmetric solvation acts as a regulatory mechanism, linking local dielectric properties to protonation energetics and motor function.

In dehydrated environments, the low dielectric constant stabilizes the protonated form of D22 and significantly elevates its p*K*_a_, thereby increasing the barrier to deprotonation. Under extreme low-dielectric conditions, deprotonation becomes energetically prohibitive, promoting proton retention and enhancing proton capture when the channel opens.

To support this mechanism, we performed FEP calculations on a model aspartate residue in both aqueous and vacuum environments, yielding a ΔΔ*G* of 76.32 kcal/mol and a corresponding p*K*_a_ shift of 53.80 (Fig. S20). Although this value exceeds typical protein p*K*_a_ ranges (which generally falls between 3 and 8), it is consistent with fundamental physical chemistry principles: in low-dielectric or gas-phase environments, deprotonation incurs a large desolvation penalty, leading to dramatically increased p*K*_a_ values.^72^

As rotation proceeds, increasing hydration around D22 lowers the local p*K*_a_ and facilitates proton release. The highly favorable free energy of proton solvation (approximately –40 to –88 kcal/mol, Table S6)^73,74^ compensates for the deprotonation barrier, enabling efficient proton transfer. Thus, water molecules actively regulate the protonation cycle rather than serving as passive solvent.

### 4.3 Hydrogen-Bonding Networks: Structural Priming for Proton Transfer

Analysis of the hydrogen-bonding networks identifies a stable proton-conduction pathway associated with F-D22 in the unplugged state (Fig. S21). The well-organized and persistent hydrogen-bonding network surrounding F-chain D22 likely serves as a preconfigured route for proton transfer. Although F-D22 is predicted to remain predominantly protonated due to its elevated p*K*_a_, the existence of this network suggests that, upon periodic changes in the local microenvironment (e.g., hydration dynamics), incoming protons from the channel can be efficiently released and relayed. This process, facilitated by the protonation state of F-D22, may proceed through a Grotthuss-like hopping mechanism.^75^ Collectively, these observations indicate that the hydrogen-bonding network around F-D22 functions as a critical structural “priming” element for the subsequent proton-transfer step in the rotational cycle.

Together, these features support a coordinated mechanism in which hydration dynamics, protonation, and structural organization are tightly coupled.

### 4.4 D22 Conformational Dynamics and Interactions with T189 and L158

Our analysis of standard MD simulations reveals that changes in the side-chain *χ*_1_ dihedral angle of D22 are tightly coupled to its protonation state, representing a key structural link between chemical state and mechanical motion (Fig. S22-23). Specifically, when a D22 site is protonated, its *χ*_1_ angle populates both *gauche* and *trans* conformations, exhibiting dynamic conformational switching (if both D22 sites are protonated, one of them will exhibit this characteristic dynamic behavior). In contrast, when D22 is deprotonated, it adopts only one fixed conformation (*gauche* for the F-D22 and *trans* for the G-D22) with no conformational fluctuation observed. The dynamic changes in the side-chain dihedral angle conformation precisely indicate that protonation triggers rotation by modulating the interaction pattern

between MotB D22 and its surrounding residues.

The conformational dynamics of D22 are closely coupled to its interaction patterns with MotA. In the deprotonated state, D22 engages primarily in electrostatic interactions, while protonation promotes hydrogen bonding with the polar residue T189. Within the dehydrated environment, the hydrogen-bonding interaction between protonated G-D22 and E-T189 is highly stable, yet thermal fluctuations can transiently break this interaction to allow minor rotation.

These disruptions coincide with a dynamic conformational change in the D22 side-chain dihedral angle, shifting from *trans* to *gauche*. Conversely, upon protonation of F-D22, it establishes new hydrogen bonds with neighboring C-T189, and its side-chain dihedral conformation switches from *gauche* to *trans*. This effectively completes the functional interchange with G-D22, supporting the stepwise rotational mechanism.

Evolutionary analysis further reveals that residue L158 plays a complementary role by maintaining a hydrophobic environment that supports D22 dehydration and proper orientation (Fig. S14, S27, 6F). Although not strictly conserved in sequence, its hydrophobic character is strongly preserved, highlighting its functional importance.

### 4.5 Previous Rotational Models

The cryo-EM structures of CjMotAB reported by Santiveri et al.^10^ established a structural basis for subsequent mechanistic investigations. Building on this framework, their study proposed that the two MotB D22 residues undergo alternating cycles of protonation and deprotonation, and highlighted the roles of key residues such as MotA T189, L158, and F186 in forming interaction interfaces and putative proton-transfer pathways.

Our results are broadly consistent with this structural framework. In addition, by utilizing refined cryo-EM structural models, we achieved improved resolution and were able to resolve several solvent molecules in the vicinity of D22. These observations suggest that water molecules play an essential role in the functional mechanism of the complex.

Through CpHMD simulations combined with free energy calculations, this study provides direct quantitative evidence—via a pronounced p*K*_a_ upshift—and a thermodynamic basis supporting the “alternating D22 protonation” hypothesis. Moreover, our analysis of hydration dynamics further substantiates the functional importance of solvent molecules, demonstrating how local water occupancy regulates the protonation equilibrium of D22.

Kubo et al. investigated the rotational mechanism using the cryo-EM structure reported by Santiveri et al., employing fixed-charge CG MD simulations.^31^ Their model suggests that maximal rotational activity occurs when both MotB D22 residues are protonated and three out of five MotA E151 residues undergo coordinated protonation switching. Based on these results, MotA E151 was proposed to function as a key “proton carrier” comparable in importance to D22.

In contrast, our all-atom CpHMD simulations based on refined cryo-EM structures do not show a significant p*K*_a_ upshift for MotA E151 under physiological conditions, indicating that these residues are unlikely to be readily protonatable in the conformations examined here. This discrepancy likely arises from differences in both methodology and structural representation. Notably, the model used by Kubo et al. lacks the N-terminal tail of MotB (residues 1–14), omitting residues such as the negatively charged MotB E14. This omission might alter the local electrostatic environment and therefore influence the predicted protonation behavior and interaction network of MotA E151.

More importantly, although CG models efficiently capture large-scale conformational motions, they inherently lack the ability to describe protonation-linked charge fluctuations, detailed electrostatic interactions, and explicit water-mediated effects. These aspects are central to the chemo-mechanical coupling mechanisms identified in our study that link D22 protonation to motor rotation. Accordingly, the functional role of MotA E151 remains an open question requiring further experimental and high-resolution computational validation. Theoretical frameworks proposed by Tu and co-workers^76–78^ describe the motor as two coupled rotating nanorings (MotA and FliG), in which rotation proceeds in discrete steps of *π/*5 (36°). In this model, alternating proton binding and release events at the two MotB D22 sites drive rotation by modulating the free energy landscape and converting the proton electrochemical gradient into mechanical torque.

Our findings provide an atomistic structural and chemical basis for this theoretical description. First, we demonstrate that the presence of at least one protonated D22 residue is a prerequisite for rotational activity, in agreement with the alternating protonation mechanism. Second, we uncover the coupling between D22 protonation states and local hydration, dielectric environment, and side-chain conformations. Together, these factors define the energetic and structural landscape in which proton binding and dissociation occur in a realistic protein–membrane environment.

Accordingly, the present work can be viewed as an atomistic realization and refinement of the theoretical model, providing detailed insight into the structural and solvation-dependent principles that govern alternating protonation.

### 4.6 Unified Mechanistic Model of CjMotAB Rotation

In summary, this study employs high-resolution all-atom simulations to directly interrogate protonation equilibria—the core process underlying the system. In doing so, we elucidate the detailed chemo-mechanical coupling that drives CjMotAB rotation. Building on these atomistic insights, we integrate the findings of prior studies to propose a comprehensive mechanistic model encompassing proton capture, hydration regulation, conformational transitions, and mechanical rotation (Fig. 7).

**Figure 7:**
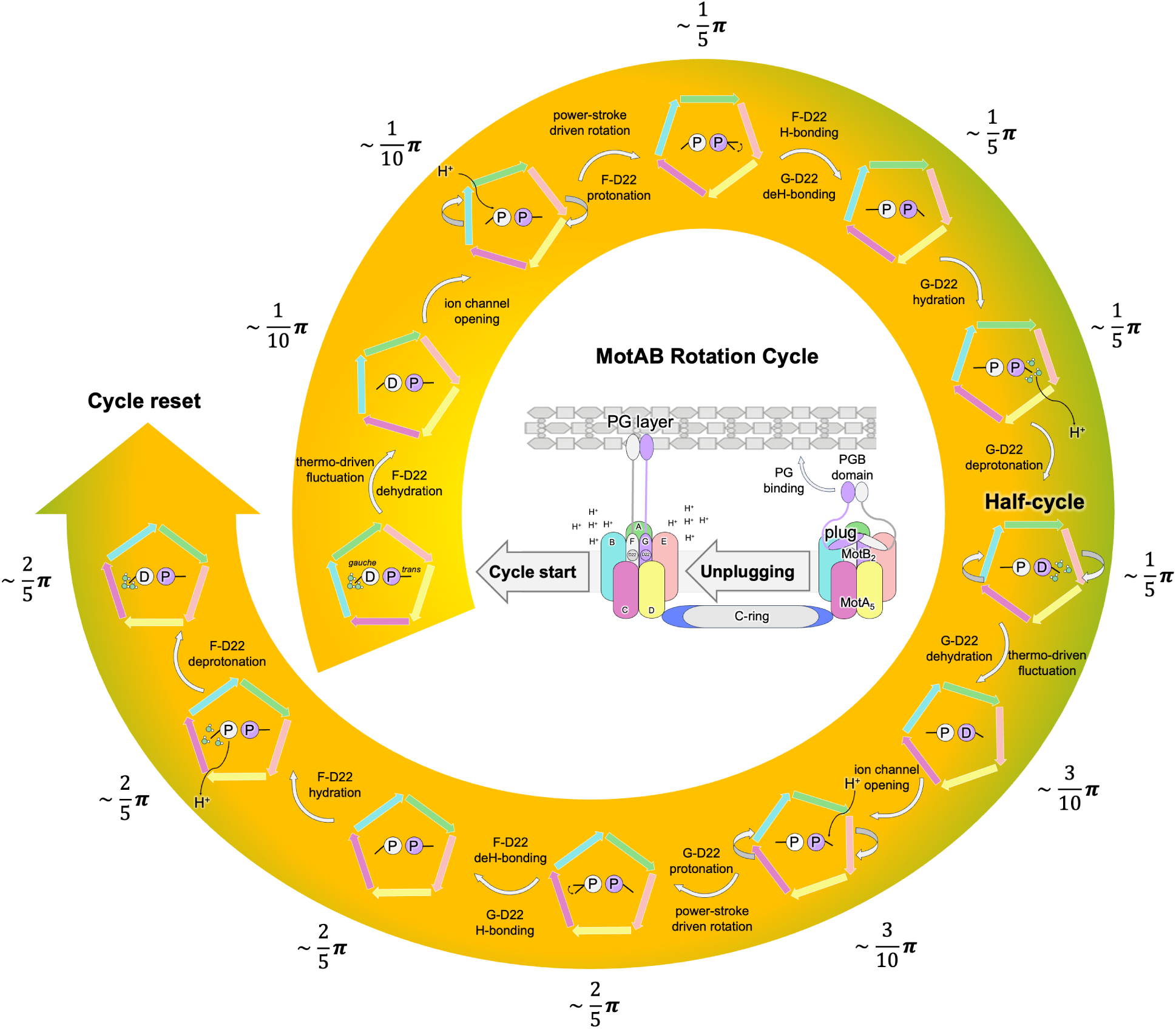
Unified mechanistic model of CjMotAB rotational cycle driven by D22 protonation, hydration dynamics, and conformational transitions. The diagram illustrates the stepwise process of CjMotAB rotation. Key features depicted include the asymmetric hydration and protonation states of F-D22 and G-D22, conformational switches of the D22 side-chain *χ*_1_ dihedral angle between *gauche* and *trans* conformations.

This unified model reconciles structural, theoretical, and simulation-based perspectives, and highlights water-mediated regulation of protonation as a central element of the mechanism. The steps of the cycle are outlined as follows:

#### • Plugging and Inactivation (Pre-active State)

The CjMotAB complex begins in a plugged, inactive configuration in which the MotB plug domain occludes the transmembrane channel. In this state, the two MotB D22 sites exhibit pronounced functional and environmental asymmetry: F-D22 is deprotonated, highly hydrated, and adopts a *gauche* conformation, whereas G-D22 is protonated, relatively dehydrated, and adopts a *trans* conformation.

#### • Unplugging, Activation, and Rotation Initiation

The initiation of rotation requires association of MotAB with the C-ring rotor to ensure effective utilization of the proton motive force (PMF) and to prevent non-productive rotation. Binding to the C-ring induces anchoring of the MotB PGB domain to the cell wall, which in turn drives displacement of the plug domain and transitions the system into the unplugged state.

Following unplugging, thermal fluctuations permit an initial small-scale rotation of MotA (approximately *π/*10). This movement weakens the hydrogen bond between G-D22 and MotA T189. Meanwhile, the hydration level around F-D22 gradually decreases, creating a low-dielectric, dehydrated microenvironment and resulting in a periplasm-facing conformation that favors proton uptake. Conformational rearrangement of the MotA F186 side chain promotes transient opening of the ion channel, enabling ion flux from the periplasm and proton uptake by F-D22.

#### • D22 Protonation-Driven Rotation

Upon protonation, F-D22 forms a new hydrogen bond with the subsequent MotA T189 residue, generating a power stroke that drives MotA through a major rotational step of *π/*5. During this process, the F-D22 side chain undergoes a *gauche-to-trans* conformational transition coupled with the rotation. The newly formed F-D22–C-T189 hydrogen bond further pulls MotB, which helps disrupt the existing G-D22–E-T189 hydrogen bond, thereby releasing the constraint on MotA and inducing a *trans-togauche* conformational switch in G-D22.

#### • Proton Release and Transport

As MotA rotates to approximately 3*π/*10, conformational rearrangements shift G- D22 into a cytoplasm-facing orientation, exposing its binding site to the cytosol and promoting rapid hydration. The resulting increase in local water density markedly lowers the p*K*_a_ of G-D22, making deprotonation thermodynamically favorable. The released proton is subsequently transported away from the site via a stable hydrogenbond network identified in this work, proceeding through a Grotthuss-like mechanism. The free energy released upon proton hydration compensates for the deprotonation barrier, completing the proton transfer process. At this stage, G-D22 stabilizes in the *gauche* conformation.

#### • D22 Role Interchange and Cycle Reset

After a single *π/*5 rotation, the local environments and functional roles of the F- and G-chain D22 residues are effectively exchanged: F-D22 becomes protonated, dehydrated, and locked in interaction with MotA in a *trans* conformation, while G-D22 becomes deprotonated, solvent-exposed, and adopts a *gauche* conformation.

The subsequent rotational step proceeds in a similar manner with the roles reversed. Thermal fluctuations allow MotA to continue minor rotating, while G-D22 begins to dehydrate in preparation for protonation. As the rotation advances to 2*π/*5, ion channel opens and G-D22 becomes protonated in the low-hydration environment and forms a hydrogen bond with the next MotA T189 residue, producing a new power stroke. Simultaneously, the F-D22 hydrogen bond is disrupted, followed by rehydration and deprotonation. The system thereby returns to a state that is chemically analogous to the initial configuration but rotated by 72°, completing a two-step, dual-proton cycle.

This model describes the full conversion of chemical energy stored in the proton electrochemical gradient into mechanical work. Proton influx first alters the protonation state of D22. Protonation states then determine side-chain conformations and hydrogen-bonding interactions, which drive periodic association and dissociation with MotA, ultimately producing stepwise directional rotation.

## 5 Conclusion

In conclusion, this study systematically investigated the protonation and conformational dynamics of titratable residues and other key residues in CjMotAB, as well as their roles in the rotational mechanism of the bacterial flagellar motor, using a combination of static structure-based p*K*_a_ prediction tools, MD simulations, and coevolutionary analysis.

Our findings include: (1) High-resolution refinement of the cryo-EM structures of CjMotAB in both plugged and unplugged states, enabling unambiguous modeling of the previously unresolved MotB N-terminal tail and key side-chain environments around the proton-carrying residue D22. (2) Pronounced and asymmetric p*K*_a_ upshifts for the two MotB D22 residues, as consistently revealed by constant-pH MD simulations and further validated by FEP calculations, confirming their role as proton carriers and establishing a thermodynamic basis for proton retention under physiological conditions. (3) Identification of two strict pre-requisites for productive MotA rotation: removal of the plug domain and protonation of at least one D22 subunit, demonstrating that unplugging alone is insufficient to drive rotation without coupled protonation dynamics. (4) Discovery of asymmetric hydration patterns surrounding F-D22 and G-D22, in which dehydration stabilizes protonation and elevated p*K*_a_ values, while rotation-coupled rehydration facilitates proton release, revealing water as an active regulator of the chemo-mechanical cycle. (5) Coupling between D22 protonation and side-chain *χ*_1_ dihedral transitions between *gauche* and *trans* conformations, which directly modulate hydrogen-bonding interactions with MotA T189 and enable directional conformational switching during rotation. (6) Evolutionarily conserved hydrophobic packing provided by MotA L158 and neighboring residues, which supports the local environment required for D22 dehydration, protonation, and productive intersubunit interactions.

Finally, we proposed an integrated mechanistic model that reconciles structural, thermodynamic, and dynamical insights from previous studies with the simulation results presented here. This model illustrates how protonation, solvation dynamics, conformational transitions, and residue-level interactions act in concert to drive the periodic rotation of CjMotAB. Central to this process is D22, which functions as a chemical switch that couples proton flux to mechanical motion, ensuring tightly coordinated and efficient energy transduction throughout the rotational cycle. Collectively, our results advance the current understanding of the CjMotAB rotational mechanism, highlighting the critical roles of D22 protonation, hydration dynamics, and conformational transitions, and providing a solid foundation for future studies on bacterial flagellar motor function and regulation.

## Acknowledgement

This work was supported by grants from the National Science Foundation of China (32371300 to Y.W., and 12174337 to P.X.). We thank Junhua Yuan for valuable discussions. Y.W. acknowledges the computational support of the Information Technology Center and State Key Lab of CAD&CG at Zhejiang University. The Novo Nordisk Foundation Center for Protein Research is supported financially by the Novo Nordisk Foundation (NNF14CC0001). N.M.I.T. acknowledges support from an NNF Hallas-Møller Ascending Investigator grant.

## Supporting Information Available

## TOC Graphic

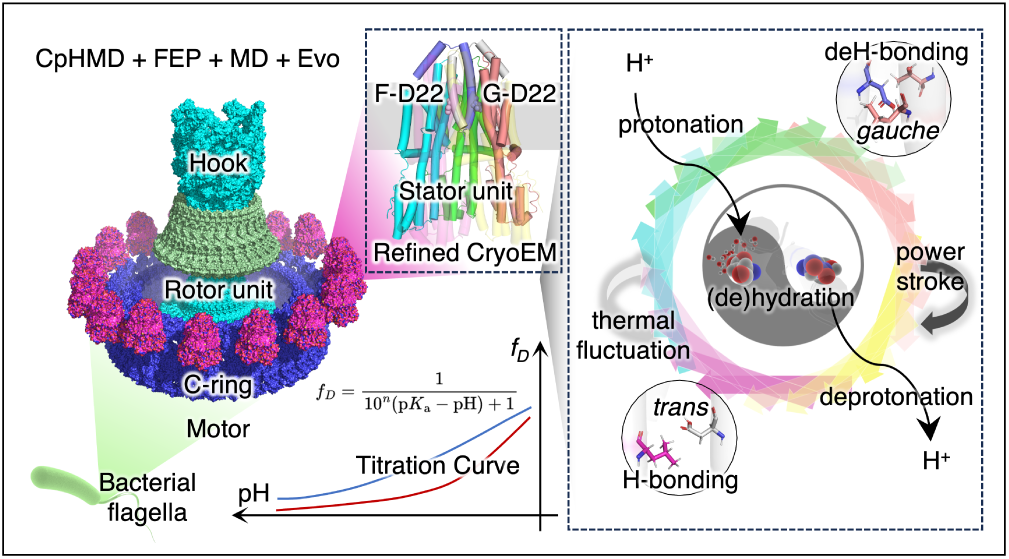

