## Supporting Information for "Asymmetric Hydration and Protonation Switching of Dual Aspartates Drive Flagellar Rotation"

### 1 Optimal Workflow for Membrane Protein CpHMD Simulation

In previous GROMACS-based CpHMD simulations of membrane proteins, simulation parameters have typically been adapted from those used for soluble proteins—a practice also implemented in our initial simulations. In the current study, we employed membrane protein-specific parameters generated via CHARMM-GUI and executed a six-step equilibration protocol. Consistent results were observed across 100 ns simulations, with no significant discrepancies identified.

Throughout both equilibration and production simulations, the temperature was maintained at 310 K using a v-rescale thermostat with a time constant of 1.0 ps. The double-well biasing potential was set to 5 kJ/mol. Based on empirical observations, we recommend this optimized workflow, which incorporates membrane protein-tailored parameters, for future CpHMD studies of membrane proteins.

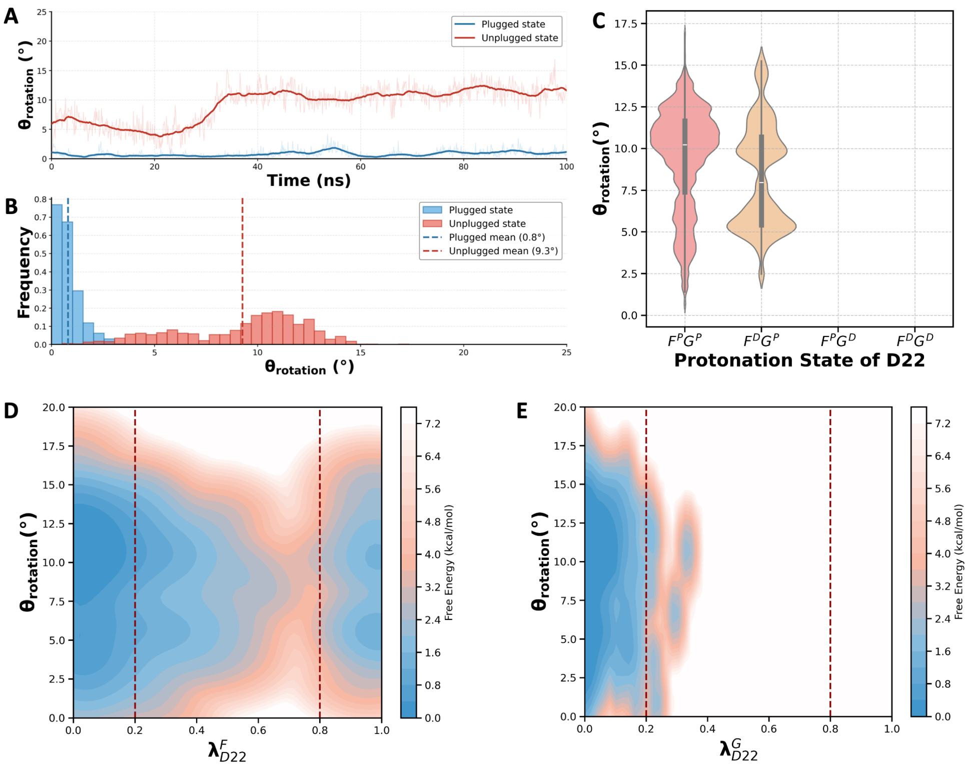


Figure S1: **CpHMD simulation results obtained with a six-step equilibration workflow.** (A) Time evolution of the MotA rotation angle ($\theta_{rotation}$) for the plugged and unplugged MotAB conformations. (B) Corresponding frequency distributions of MotA rotation angles in the plugged and unplugged states. (C) Dependence of the rotation angle on the combined protonation states of the F and G chains. Here, 'P' denotes a protonated state and 'D' denotes a deprotonated state; the first character corresponds to the F chain, and the second to the G chain. (D, E) Free energy surfaces as a function of the MotA rotation angle relative to MotB ($\theta_{rotation}$) and the protonation coordinate of D22 ($\lambda_{D22}$), derived from CpHMD simulations of the unplugged MotAB state.

#### 2 Cryo-EM Structure of CjMotAB

Herein, we present the relevant results and methodological details of the cryo-EM experiments conducted to resolve the CjMotAB structure.

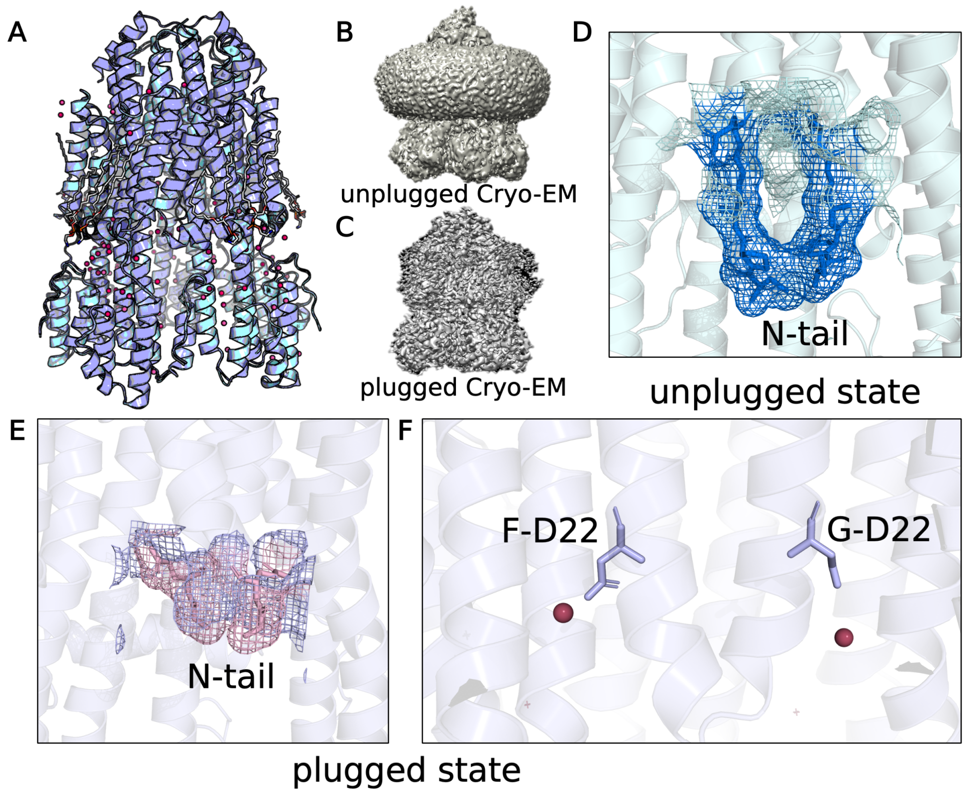


Figure S2: Structural characterization of CjMotAB in unplugged and plugged conformational states. (A) Structural superposition of the full-length CjMotAB complexes in the unplugged (cyan) and plugged (purple) states. (B, C) cryo-EM density maps of the CjMotAB complex in the (B) unplugged and (C) plugged states, visualizing the distinct overall architectures. (D, E) Cryo-EM density maps (blue mesh for unplugged, pink mesh for plugged) and fitted atomic models of the N-terminal tail (N-tail) region, demonstrating the structural differences between the two states. (F) Cryo-EM density and atomic model of the conserved residue D22 (F-D22 and G-D22) in the plugged state, with the coordinated water molecules (red spheres) shown in the binding pocket.

Table S1: Cryo-EM data collection, refinement and model statistics.

| | **CjMotAB WT** | **CjMotAB_unplugged** |
|---|---|---|
| **Data collection and processing** | | |
| Microscope | Titan Krios G2 | Titan Krios G2 |
| Magnification (nominal) | 96,000× | 96,000× |
| Voltage (kV) | 300 | 300 |
| Total exposure ($e^-/Å^2$) | 40.84 | 42.51 |
| Exposure fractions (no.) | 40 | 40 |
| Pixel size (Å) | 0.832 | 0.832 |
| Symmetry imposed | C1 | C1 |
| Movies used (no.) | 2,838 | 5,109 |
| Final particles (no.) | 230,686 | 446,432 |
| Box size (pixels) | 400 | 400 |
| Map resolution (Å) (FSC 0.143) | 2.67 | 2.29 |
| **Refinement** | | |
| Refinement resolution (Å) (FSC map vs. model (masked)=0.143) | 2.60 | 2.29 |
| **Model composition** | | |
| Non-hydrogen atoms | 10,579 | 10,864 |
| Protein residues | 1,363 | 1,336 |
| Water molecules | 123 | 201 |
| **B-factors (mean; $Å^2$)** | | |
| Protein | 75.49 | 71.25 |
| Solvent | 63.74 | 66.82 |
| **R.m.s. deviations** | | |
| Bond lengths (Å) | 0.006 | 0.003 |
| Bond angles (°) | 0.658 | 0.521 |
| CC (mask) | 0.81 | 0.80 |
| **Validation** | | |
| MolProbity score | 1.8 | 1.43 |
| Poor rotamers (%) | 2.79 | 0.55 |
| **Ramachandran plot** | | |
| Favored (%) | 98.30 | 98.18 |
| Allowed (%) | 1.7 | 1.82 |
| Disallowed (%) | 0.00 | 0.00 |

Table S2: Summary of all simulation systems

| Model | Simulation Method | D22 Protonation State | Simulation Time per Run | Replicas | Temperature |
|---|---|---|---|---|---|
| Plugged | CpHMD | Titratable | 100 ns | 12 pH replicas × 6 | 300 K |
| Unplugged | CpHMD | Titratable | 100 ns | 12 pH replicas × 6 | 300 K |
| Unplugged | CpHMD (six-step equilibration protocol) | Titratable | 100ns | 12 pH replicas × 1 | 310K |
| Plug-removed | Standard MD | $F^DG^D$ | 1000 ns | 1 | 300 K |
| Plug-removed | Standard MD | $F^DG^P$ | 1000 ns | 1 | 300 K |
| Plug-removed | Standard MD | $F^PG^D$ | 1000 ns | 1 | 300 K |
| Unplugged | Standard MD | $F^PG^P$ | 1000 ns | 1 | 300 K |
| Unplugged | FEP | $F^P \to F^D$ | 10 ns | 11 replicas × 1 | 300 K |
| Unplugged | FEP | $G^P \to G^D$ | 15 ns | 21 replicas × 1 | 300 K |
| Aqueous Asp | FEP | $D^P \to D^D$ | 10 ns | 11 replicas × 1 | 300 K |
| Unplugged | FEP | $F^P \to F^D$ | 10 ns | 11 replicas × 1 | 310 K |
| Unplugged | FEP | $G^P \to G^D$ | 15 ns | 21 replicas × 1 | 310 K |
| Aqueous Asp | FEP | $D^P \to D^D$ | 10 ns | 11 replicas × 1 | 310 K |
| Unplugged | FEP (vdw-remained) | $F^P \to F^D$ | 10 ns | 11 replicas × 1 | 300 K |
| Unplugged | FEP (vdw-remained) | $G^P \to G^D$ | 15 ns | 21 replicas × 1 | 300 K |
| Aqueous Asp | FEP (vdw-remained) | $D^P \to D^D$ | 10 ns | 11 replicas × 1 | 300 K |
| Unplugged | FEP (vdw-remained) | $F^P \to F^D$ | 10 ns | 11 replicas × 1 | 310 K |
| Unplugged | FEP (vdw-remained) | $G^P \to G^D$ | 15 ns | 21 replicas × 1 | 310 K |
| Aqueous Asp | FEP (vdw-remained) | $D^P \to D^D$ | 10 ns | 11 replicas × 1 | 310 K |
| Vacuum Asp | FEP | $D^P \to D^D$ | 10 ns | 11 replicas × 1 | 310 K |

### 3 Titratable Curves of Important Residue Sites

To illustrate the titration behavior of functionally relevant residues, titration curves were plotted individually for clarity. Specifically, these curves correspond to residue E151 on MotA, and residues E14 and D22 on MotB—each analyzed in both unplugged and plugged conformations. All titration curves were fitted using the Henderson-Hasselbalch equation, while block analysis was employed to evaluate the standard error of the mean (SEM).

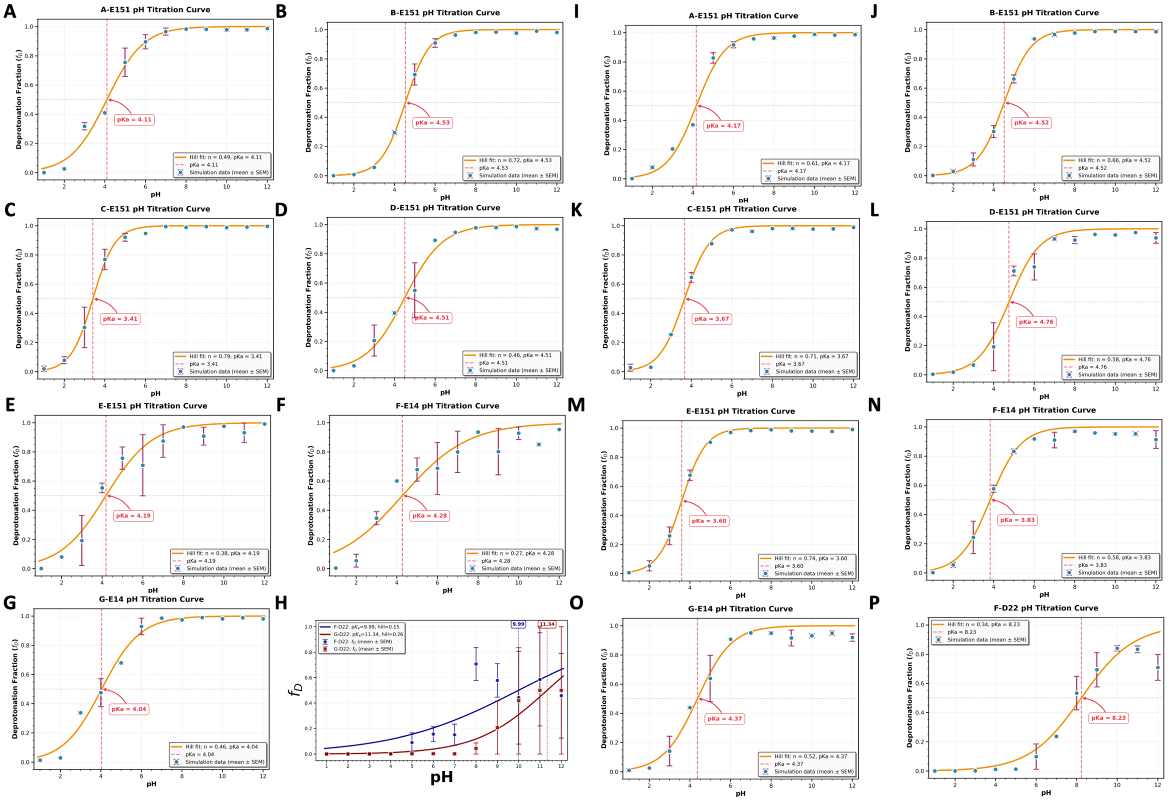


Figure S3: **pH titration curves and fitted p$K_a$ values for critical ionizable residues in the unplugged and plugged states of CjMotAB.** (A-H) Titration curves for key residues in the unplugged state, with fitted p$K_a$ values and error estimates labeled. Panel H shows the deprotonation fraction ($f_D$) as a function of pH for selected residues. (I-P) Corresponding titration curves and fitted p$K_a$ values for the same critical residues in the plugged state.

### 4 The Convergence of p$K_a$ Calculation

To assess the convergence of p$K_a$ calculations, we analyzed variations in the mean degree of deprotonation and its standard error across different pH values and block sizes.

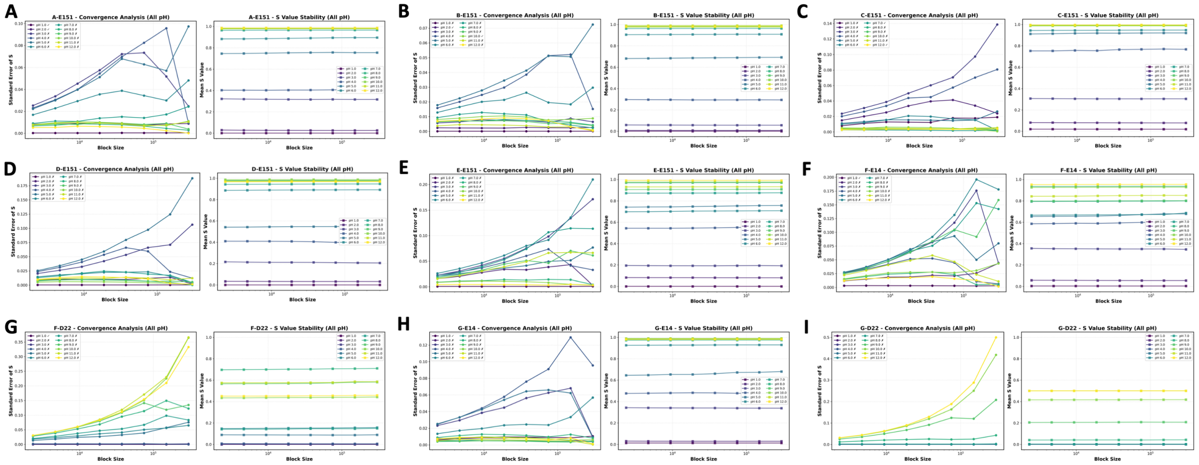

Figure S4: **Titration convergence analysis of all critical ionizable residues in the unplugged state of CjMotAB.** (A–I) For each residue, left panels show the standard error of the protonation fraction (S) as a function of block size (block averaging convergence), and right panels show the stability of the mean S value across block sizes, across all simulated pH conditions.

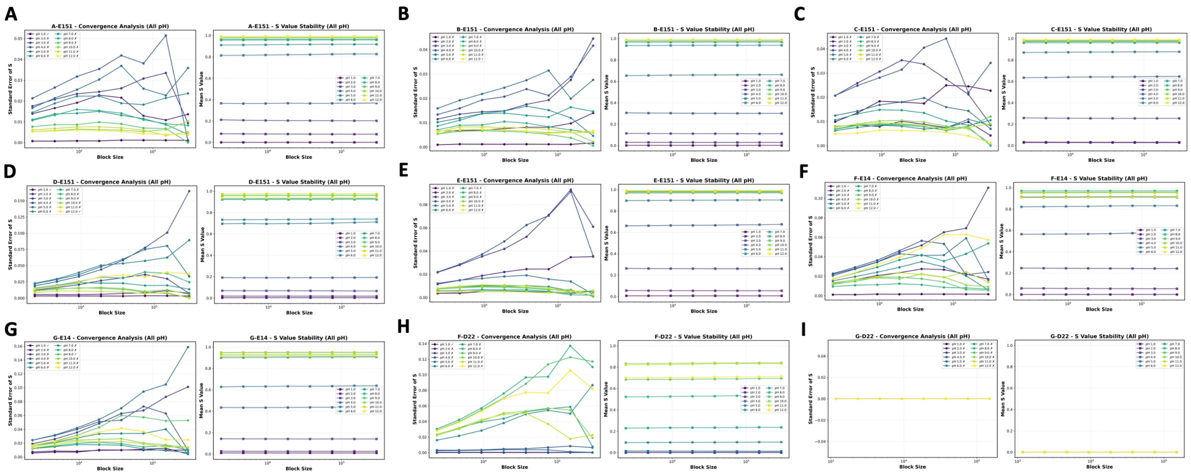


Figure S5: **Titration convergence analysis of all critical ionizable residues in the plugged state of CjMotAB.** (A–I) For each residue, left panels show the standard error of the protonation fraction (S) as a function of block size (block averaging convergence), and right panels show the stability of the mean S value across block sizes, across all simulated pH conditions.

### 5 p$K_\mathrm{a}$ Shifts of All Titratable Sites

To systematically evaluate the influence of protonation/deprotonation events across all titratable sites on the p$K_\mathrm{a}$ values of the entire system and functionally critical residues, we designated all aspartic acid (D), glutamic acid (E), and histidine (H) residues as titratable in our constant-pH molecular dynamics (CpHMD) simulations. The p$K_\mathrm{a}$ shifts of all titratable residues are shown in Fig. S6 (A–E for the unplugged state, G–L for the plugged state). However, histidine residues showed poor convergence in our CpHMD simulations, and our primary research focus centers on the functionally relevant aspartic and glutamic acid residues. Therefore, all subsequent analyses were restricted to titratable D and E residues. To ensure the reliability of our results, we calculated p$K_\mathrm{a}$ values using three complementary approaches: PropKa 3.0, DeepKa, and CpHMD, and performed cross-comparisons to validate the consistency of the predictions.

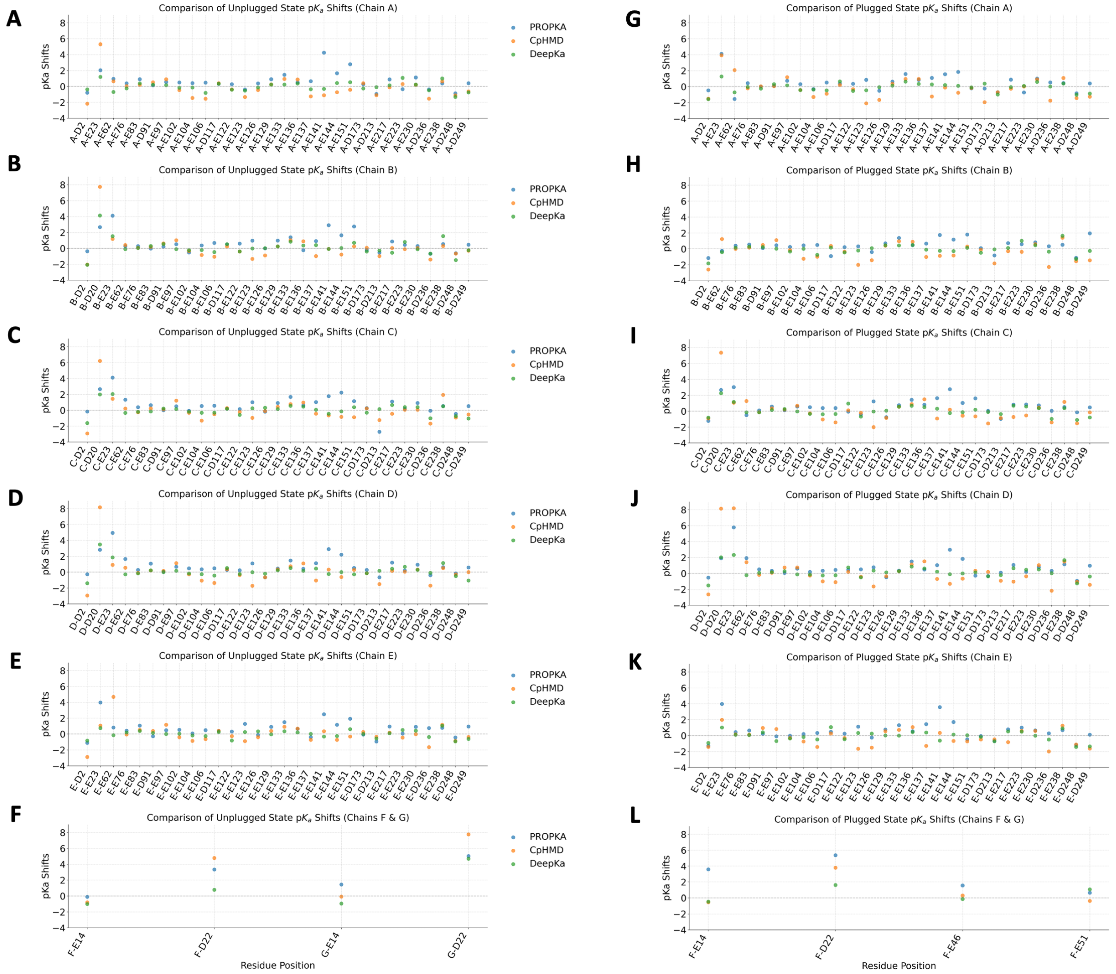


Figure S6: **Comparison of** p$K_\mathrm{a}$ **shifts for titratable sites in unplugged and plugged CjMotAB states.** (A-F) p$K_\mathrm{a}$ shifts calculated via PROPKA, CpHMD, and DeepKa for all titratable residues in the unplugged state, across Chains A–F (F includes Chains F and G). (G-L) p$K_\mathrm{a}$ shifts calculated via PROPKA, CpHMD, and DeepKa for all titratable residues in the plugged state, across Chains A–F (F includes Chains F and G).

### 6 Comparison Among Three p$K_a$ Prediction Methods

To assess the consistency and reliability of p$K_a$ shift predictions across different computational approaches, we performed pairwise correlation analyses of the calculated values. Notably, CpHMD, PropKa 3.0, and DeepKa all exhibited strong correlations (Pearson correlation coefficient $R \geq 0.5$) for both the unplugged (Fig. S7) and plugged (Fig. S8) conformational states of the membrane protein. Our simulation system focuses on a membrane protein with dynamic conformational heterogeneity, where functionally critical titratable residues are localized within the transmembrane domain. However, PropKa 3.0 and DeepKa are primarily optimized for soluble protein systems, as their training datasets are overwhelmingly composed of soluble protein structures. In contrast, the CpHMD method (developed by Hess et al.) explicitly enables all-atom simulations under the NPT ensemble, rendering it inherently more suitable for membrane protein systems and their complex transmembrane environments.

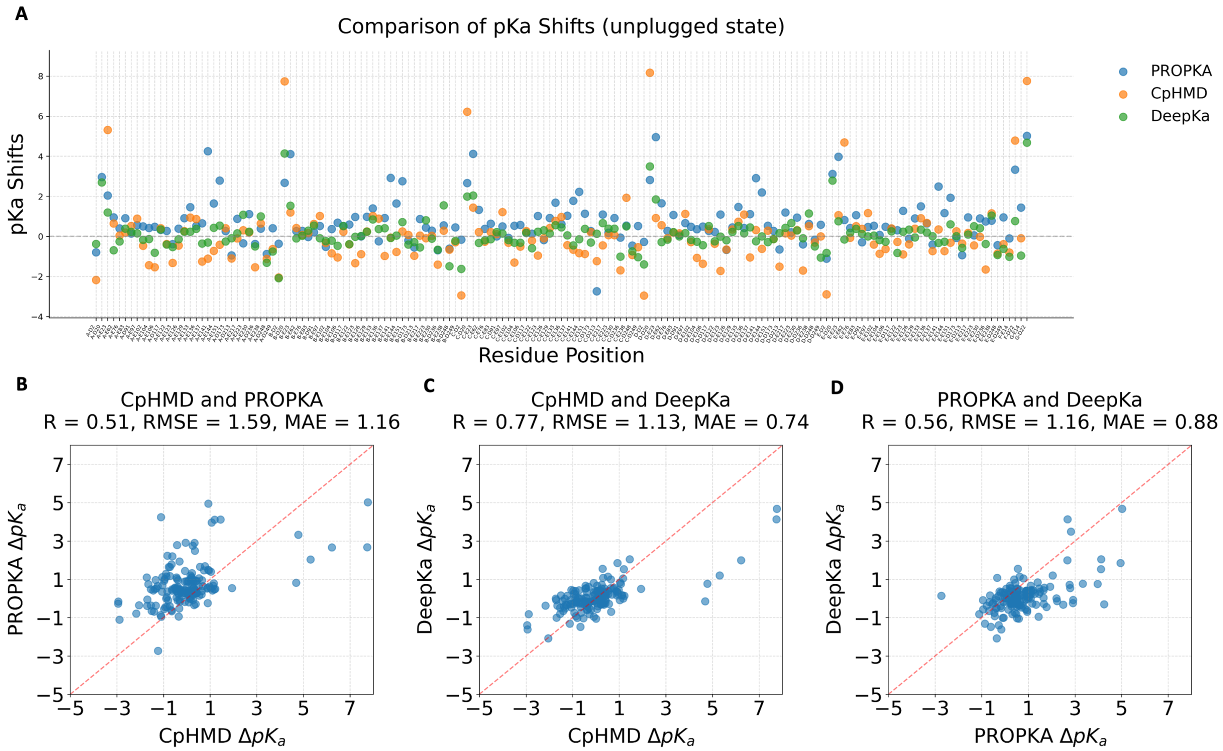


Figure S7: p$K_\mathrm{a}$ **shift analysis of titratable residues in the unplugged state of Cj-MotAB.** (A) $\Delta$p$K_\mathrm{a}$ (p$K_\mathrm{a}$ shift) values for all titratable residues, calculated by three methods: PROPKA (blue), CpHMD (orange), and DeepKa (green). (B–D) Pairwise comparisons of $\Delta$p$K_\mathrm{a}$ predictions between the three methods, with correlation coefficient (R), root-mean-square error (RMSE), and mean absolute error (MAE) reported for each comparison: (B) CpHMD vs. PROPKA, (C) CpHMD vs. DeepKa, (D) PROPKA vs. DeepKa.

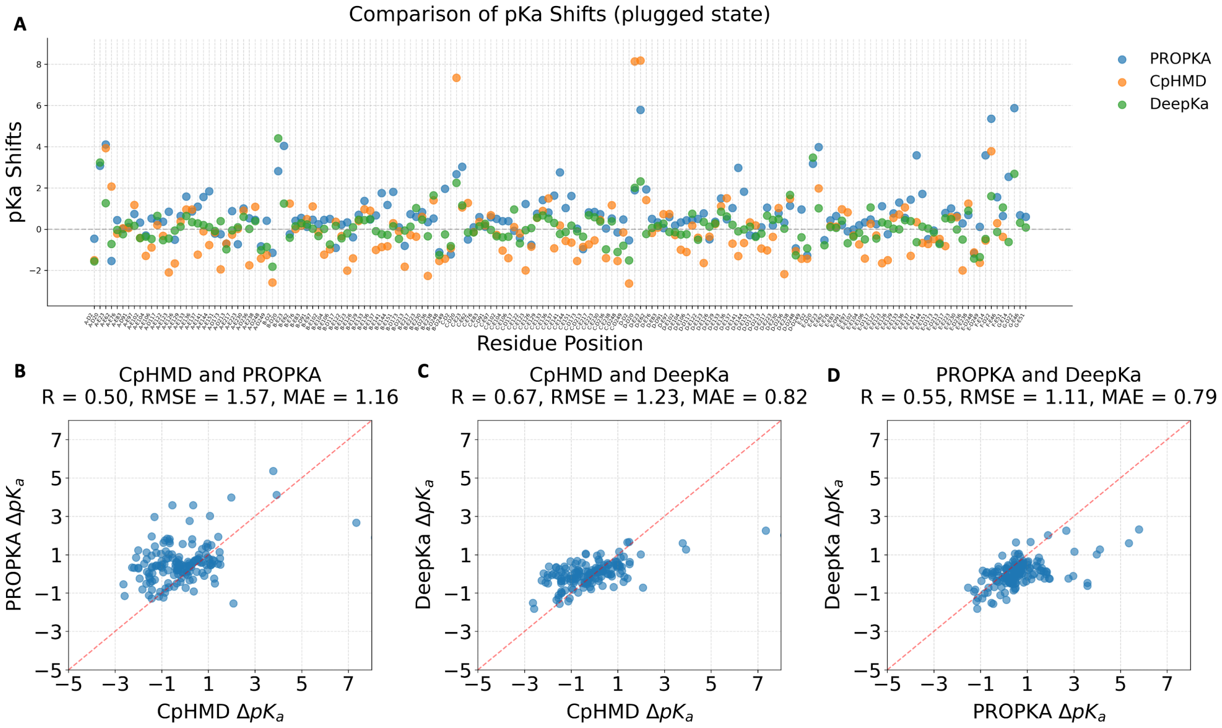


Figure S8: p$K_a$ **shift analysis of titratable residues in the plugged state of Cj-MotAB.** (A) $\Delta$p$K_a$ (p$K_a$ shift) values for all titratable residues, calculated by three methods: PROPKA (blue), CpHMD (orange), and DeepKa (green). (B–D) Pairwise comparisons of $\Delta$p$K_a$ predictions between the three methods, with correlation coefficient (R), root-mean-square error (RMSE), and mean absolute error (MAE) reported for each comparison: (B) CpHMD vs. PROPKA, (C) CpHMD vs. DeepKa, (D) PROPKA vs. DeepKa.

### 7 FEP Calculation Under 310K

Free energy differences were also calculated at 310 K, given that this temperature is more physiologically relevant for membrane protein simulations. All computational procedures were consistent with those employed for the 300 K calculations. The results are presented in Table S1.

Table S3: FEP Calculation under 310K

| 310K | F chain D22 (kcal/mol) | G chain D22 (kcal/mol) |
|---|---|---|
| $\Delta G_{ref}$ | -58.92 | |
| $\Delta G_{prot}$ | -47.74 | -23.51 |
| $\Delta\Delta G$ | 11.18 | 35.41 |
| p$K_a$ shift | 7.88 | 24.96 |

### 8 FEP Calculations With Retained Proton Van der Waals Interactions

In standard CpHMD implementations, the primary difference between protonated and deprotonated residue states is defined by their charge states, with the bound hydrogen atom retained and its Van der Waals (VdW) interactions fully preserved. While prior work has shown that this feature has negligible impact on soluble protein systems, we performed a series of FEP calculations to quantify the magnitude of such effects in membrane proteins. Our results demonstrate that VdW interactions contribute only a minor difference to the calculated free energies.

Table S4: FEP Calculation with Van der Waal Force Remained under 300K

| 300K | F chain D22 (kcal/mol) | G chain D22 (kcal/mol) |
|---|---|---|
| $\Delta G_{ref}$ | -59.41 | |
| $\Delta G_{prot}$ | -47.91 | -22.67 |
| $\Delta\Delta G$ | 11.5 | 36.74 |
| p$K_a$ shift | 8.38 | 26.76 |

Table S5: FEP Calculation with Van der Waal Force Remained under 310K

| 310K | F chain D22 (kcal/mol) | G chain D22 (kcal/mol) |
|---|---|---|
| $\Delta G_{ref}$ | -59.21 | |
| $\Delta G_{prot}$ | -49.45 | -25.36 |
| $\Delta\Delta G$ | 9.76 | 33.85 |
| p$K_a$ shift | 6.88 | 23.86 |

### 9 The Convergence of FEP Calculation

To ensure the accuracy and convergence of our FEP calculations, alchemical free energy methods were employed for the processing of simulation data. Furthermore, three complementary free energy estimation approaches—including the Multistate Bennett Acceptance Ratio (MBAR)—were utilized to compute the free energy differences.

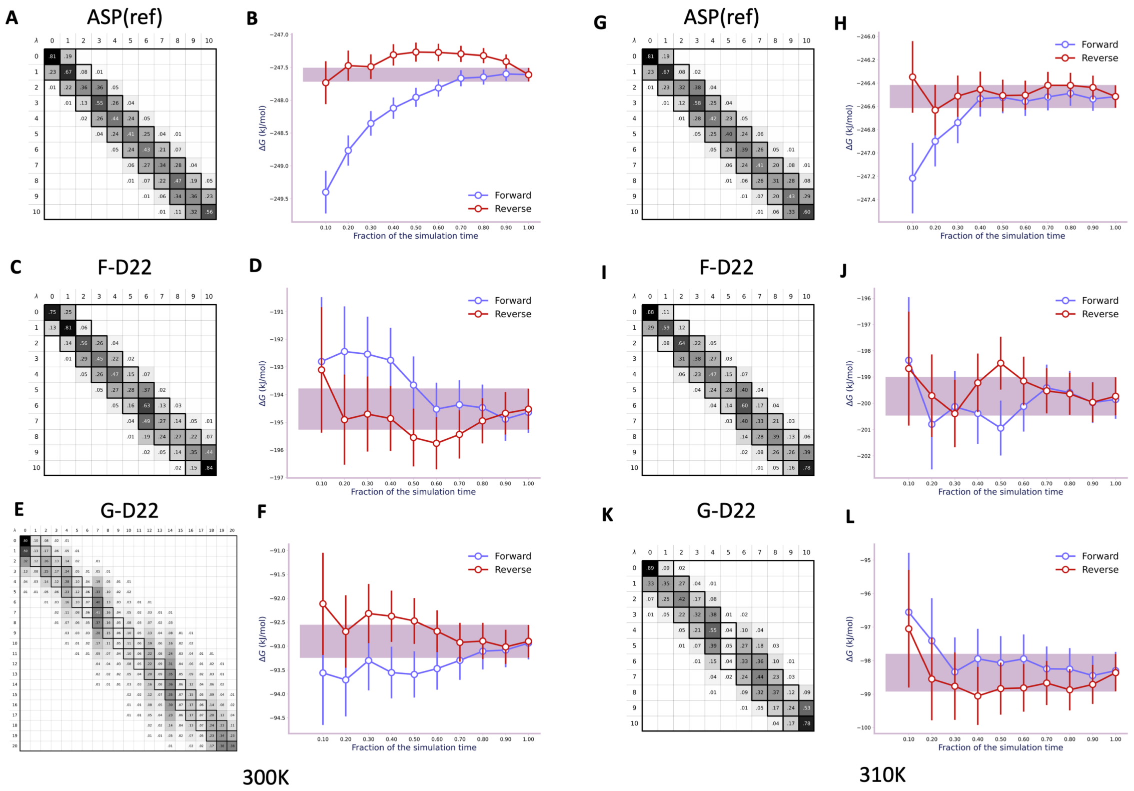


Figure S9: **Convergence and error analysis of FEP calculations at 300 K and 310 K.** (A–F) Results at 300 K: (A, C, E) Free energy matrices for reference aspartic acid (ASP(ref)), F-D22, and G-D22, respectively; (B, D, F) Corresponding forward/reverse FEP convergence profiles, with the purple shaded region indicating the converged free energy estimate. (G–L) Results at 310 K: (G, I, K) Free energy matrices for ASP(ref), F-D22, and G-D22, respectively; (H, J, L) Corresponding forward/reverse FEP convergence profiles, with the purple shaded region indicating the converged free energy estimate.

To clarify the role of D22 protonation in the rotational mechanism, we categorized all simulated conformations based on the protonation states of the D22 residues in the F and G chains of CpHMD simulations. This classification yields four distinct states: $F^P G^P$, $F^D G^P$,

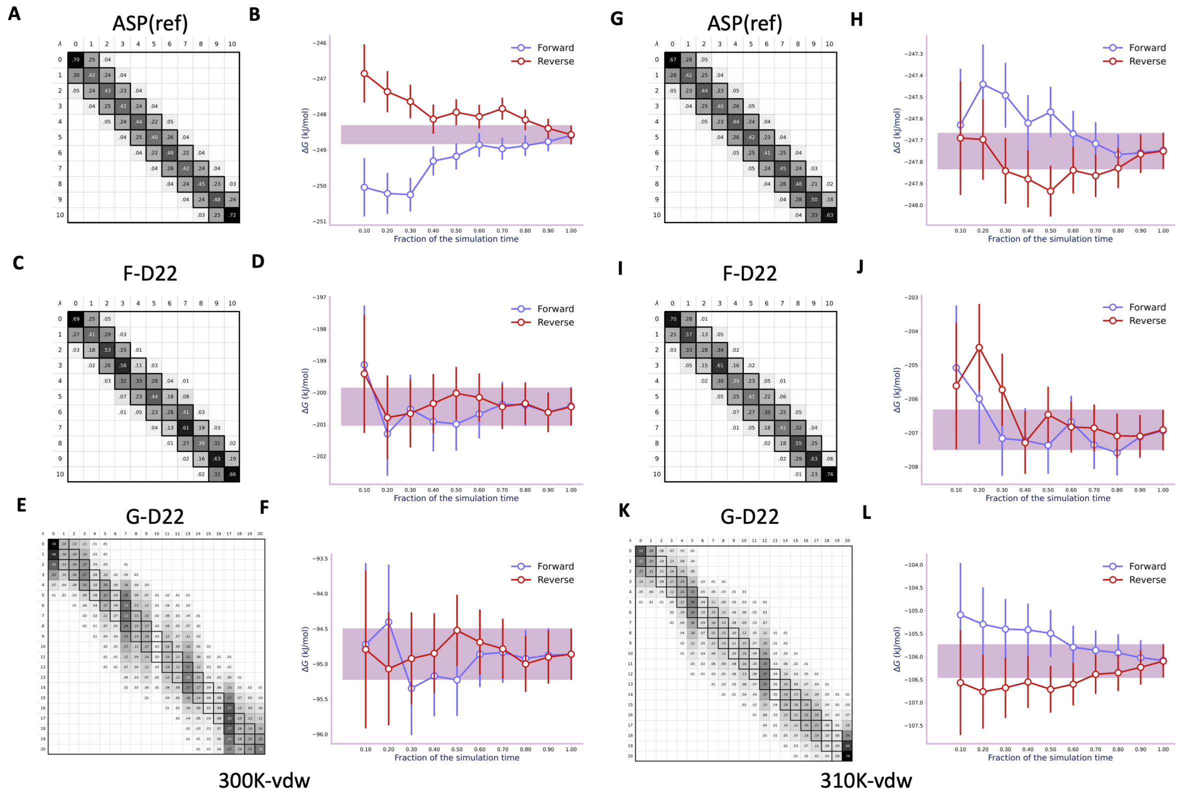


Figure S10: **Convergence and error analysis of FEP calculations with retained proton Van der Waals (vdW) interactions, performed at 300 K and 310 K.** (A–F) Results at 300 K (300K-vdw): (A, C, E) Free energy matrices for reference aspartic acid (ASP(ref)), F-D22, and G-D22, respectively; (B, D, F) Corresponding forward/reverse FEP convergence profiles, with the purple shaded region indicating the converged free energy estimate. (G–L) Results at 310 K (310K-vdw): (G, I, K) Free energy matrices for ASP(ref), F-D22, and G-D22, respectively; (H, J, L) Corresponding forward/reverse FEP convergence profiles, with the purple shaded region indicating the converged free energy estimate.

$F^PG^D$, and $F^DG^D$, where the superscripts P and D denote protonated and deprotonated states, respectively. As illustrated in Fig. S11, the horizontal axis represents the four possible combinations of D22 protonation states, while the vertical axis indicates the rotation angle. Notably, all conformations containing at least one protonated D22 residue exhibited rotational motion.

We also calculated free energy surfaces as a function of the protonation states of D22 in the F and G chains ($\lambda_{\mathrm{D22}}$) and the MotA rotation angle ($\theta_{\mathrm{rotation}}$). The FES reveal that when both F-D22 and G-D22 are protonated ($\lambda^F_{\mathrm{D22}} \approx 0, \lambda^G_{\mathrm{D22}} \approx 0$), as well as when F-D22 is deprotonated ($\lambda^F_{\mathrm{D22}} \approx 1$), states characterized by a substantial MotA rotation also correspond to low-energy basins. This indicates that the rotational motion of MotA was successfully sampled across these three protonation combinations throughout the CpHMD simulations.

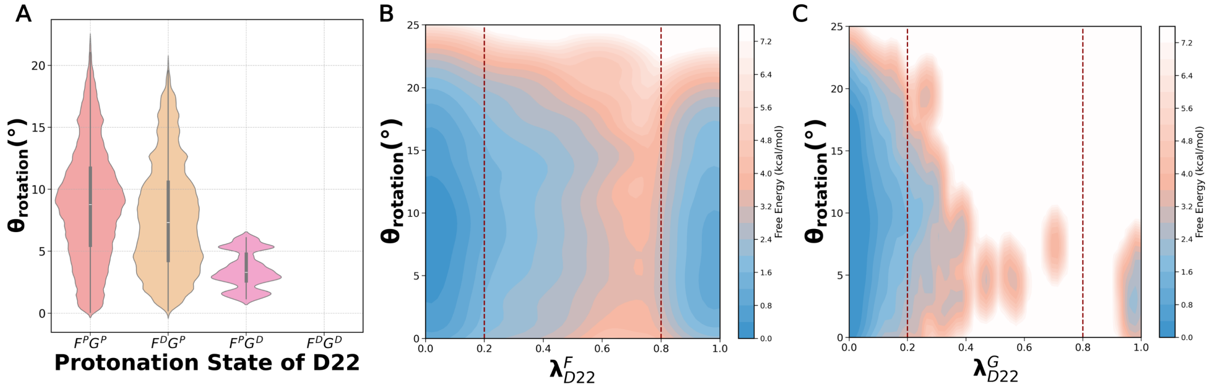


Figure S11: **Protonation-coupled conformational dynamics of the unplugged MotAB state from CpHMD simulations.** (A) Violin plots showing the distribution of MotA rotation angles ($\theta_{rotation}$) as a function of the combined protonation states of D22 residues on the F and G chains (P = protonated, D = deprotonated). (B, C) Two-dimensional free energy surfaces (FES) plotted as a function of $\theta_{rotation}$ and the protonation coordinate of D22 on the (B) F chain ($\lambda^F_{D22}$) and (C) G chain ($\lambda^G_{D22}$), illustrating the coupling between MotA rotation and residue protonation.

### 10 Hydrophobic Environment Around D22 sites

To further elucidate the elevated p$K_\mathrm{a}$ of the G-D22, the hydrophobic environments surrounding the two MotB D22 sites were analyzed. As illustrated in Fig. S12, the G-D22 is situated closer to highly hydrophobic residues (e.g., Leu, Val, Ile, and Phe) than the F-D22. Solvent-accessible surface area (SASA) analysis corroborates this trend, albeit with a modest difference between the two sites. Consequently, the G-D22 resides in a more hydrophobic environment. Collectively, these results suggest that the F-D22 may be involved in the proton-release phase, whereas the G-D22 remains protonated and drives the rotational motion. Additionally, several key hydrophobic residues were identified: L158 in chains B, C, and E; P186 in chains B and C; I193 in chains B and E; V162 in chain E; and V18, P23, L24, L26, and L27 in chains F and G.

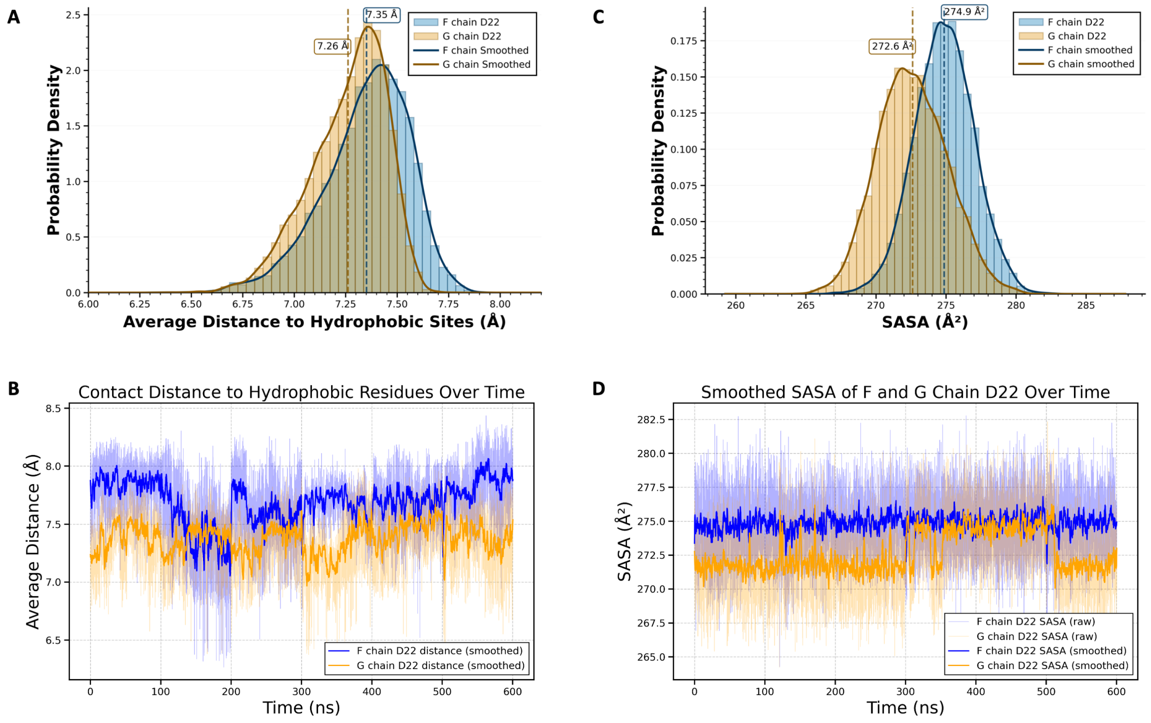


Figure S12: **Characterization of the hydrophobic environment surrounding D22 residues at pH 7 from CpHMD simulations.** (A) Probability density distributions of the average distance from D22 to hydrophobic sites for the F and G chains, with peak values indicated. (B) Time evolution of the average distance from D22 to hydrophobic residues over 600 ns of simulation. (C) Probability density distributions of the solvent-accessible surface area (SASA) of D22 for the F and G chains, with peak values indicated. (D) Time evolution of the smoothed SASA of D22 residues over 600 ns of simulation.

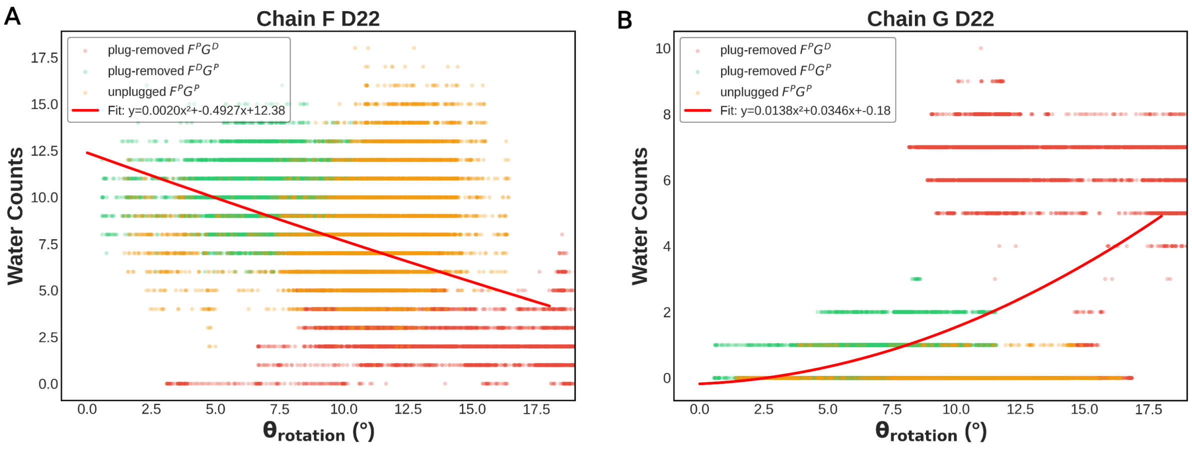


Figure S13: **Hydration of D22 residues coupled to MotA rotation angle in standard MD simulations.** (A) Scatter plot and quadratic fit showing the relationship between the MotA rotation angle ($\theta_{rotation}$) and water counts at F-D22, across different conformational states. (B) Correlation analysis for G-D22, illustrating the dependence of water counts on $\theta_{rotation}$ and protonation states.

### 11 Side-chain Dynamics and Interaction Analysis

Further contact analysis of the unplugged-state conformation of CjMotAB highlighted interactions around the D22 residues in CpHMD simulations. For MotB D22, Fig. 6A–C illustrates that D22 in the F chain is positioned between T189 in chain B and L158 in chain C, whereas D22 in the G chain is located between L158 in chain D and T189 in chain E.

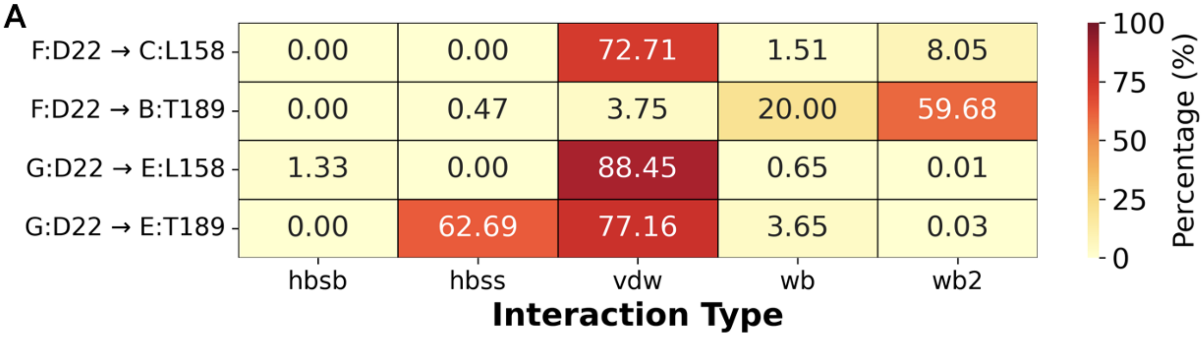


Figure S14: (A) Heatmap depicting interactions between MotB D22 and MotA residues L158 and T189. (hbsb: hydrogen bond between side chain and backbone; hbss: hydrogen bond between side chain and side chain; vdw: van der Waals force; wb: water bridge; wb2: double water molecule bridge; hbbb: hydrogen bond between backbone and backbone.

To illustrate the rotational mechanism of MotAB from the dynamics view, we analyze the contact in CpHMD simulations of the unplugged state.

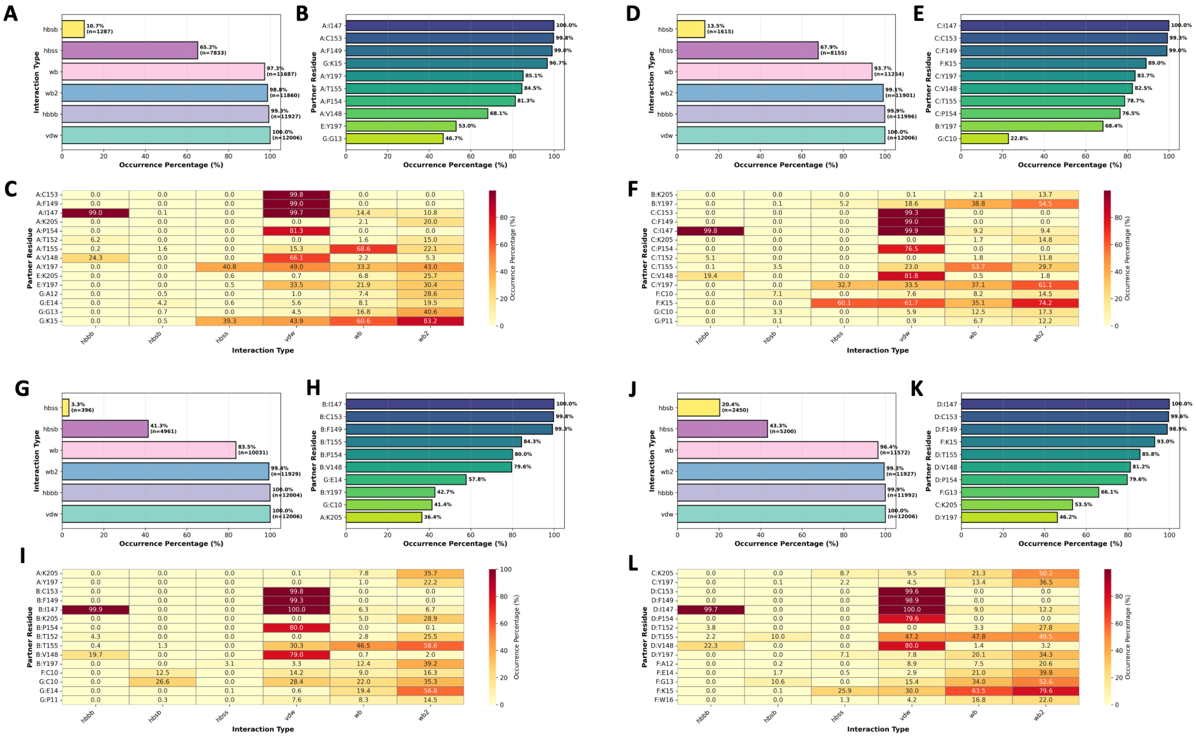


Figure S15: (A-L) Chain A-D interaction type, partner, and occurrence percentage of E151.

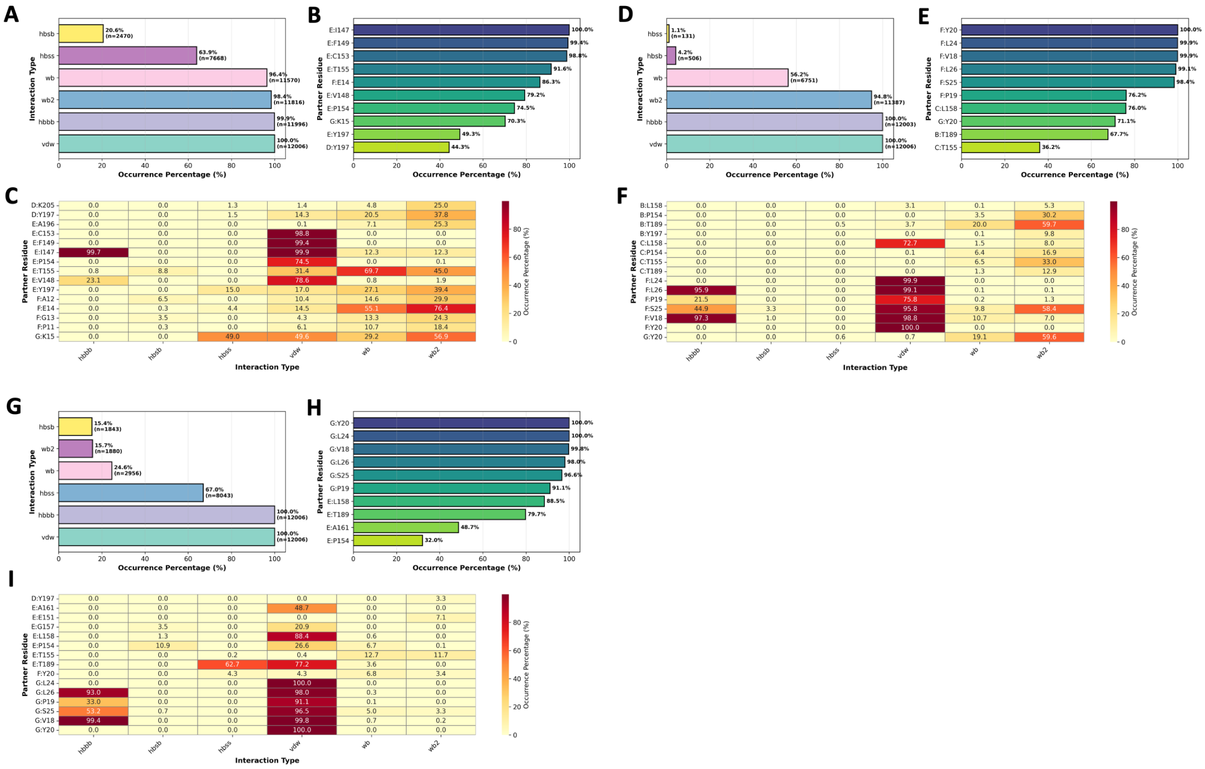


Figure S16: (A-C) Chain E interaction type, partner, and occurrence percentage of E151. (D-I) Chain F and G interaction type, partner, and occurrence percentage of D22.

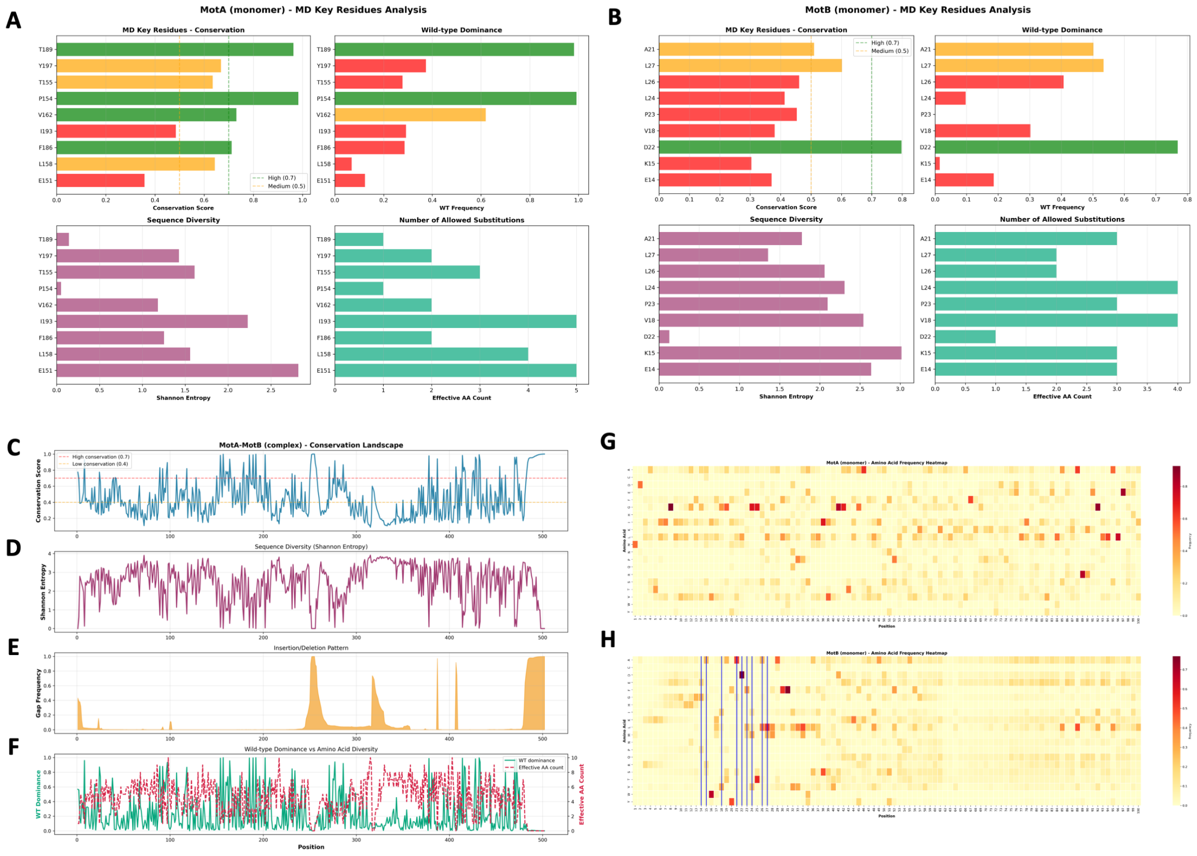


Figure S17: **Evolution Analysis of MotAB key residues.** (A-B) The sequence conservation of key residues of MotA and MotB. (C) The sequence conservation of all residues of MotA and MotB. (D) The Shannon entropy of the sequence diversity of MotAB. (E) The sequence gap analysis of MotAB. (F) The dominance of wild type and the number of allowed substitutions. (G-H) Amino acid frequency heatmap of MotAB.

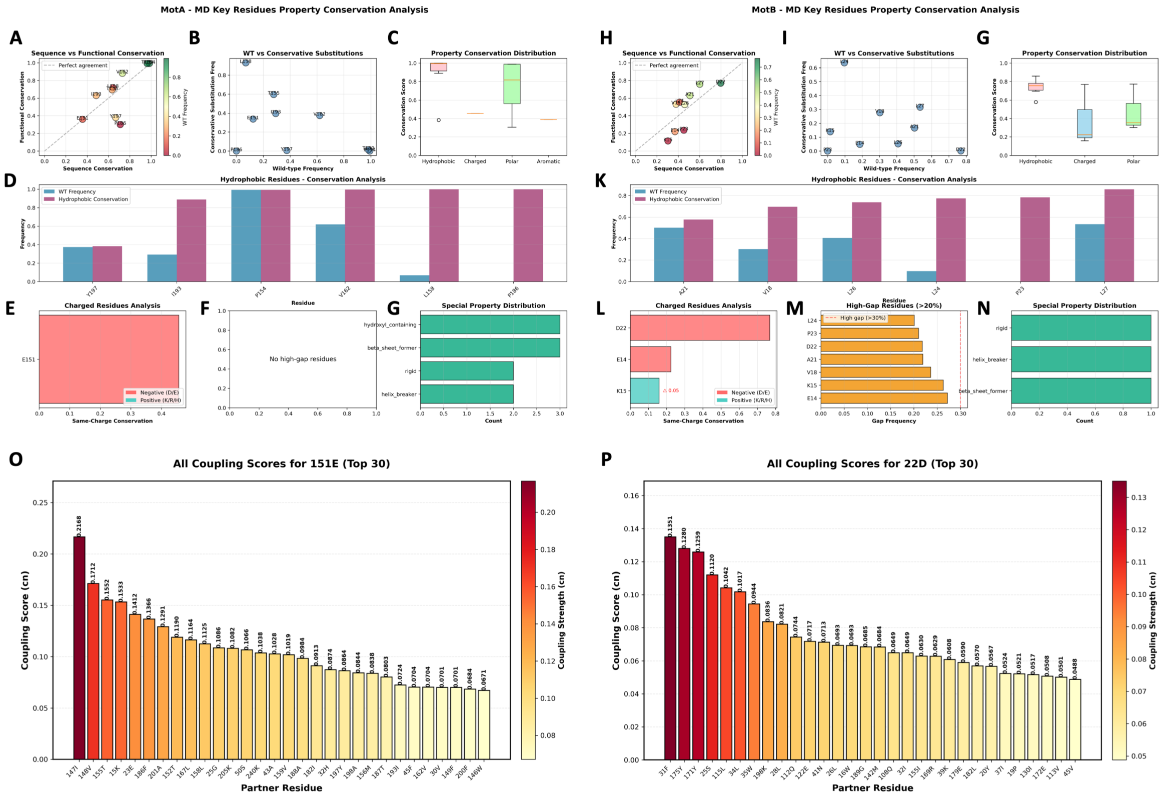


Figure S18: **Evolutionary conservation and residue coupling analysis of key functional residues in the MotAB stator complex.** (A–N) Multi-dimensional property conservation analysis for critical residues in MotA (A–G) and MotB (H–N), including: sequence vs. functional conservation, conservative substitution analysis, property conservation distribution, hydrophobic/charged residue conservation, gap frequency analysis, and special property distribution. (O–P) Top 30 residue coupling scores for (O) MotA E151 and (P) MotB D22, colored by coupling strength, highlighting their key co-evolutionary interaction partners.

In addition to the epistatic model, EVcouplings can also compute mutation sensitivity scores using an independent model that ignores inter-site couplings. By comparing the results of these two models, we found that all the residues identified in our study are evolutionarily beneficial, indicating they confer selective advantages during evolutionary processes.

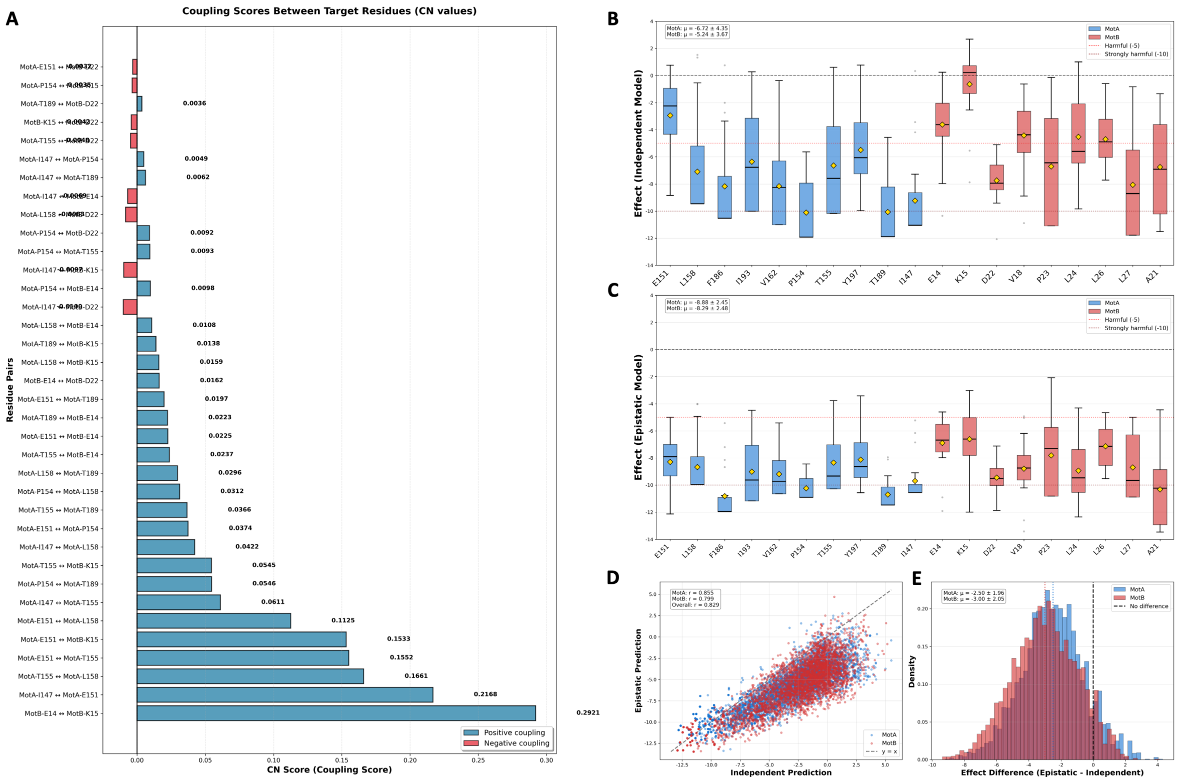


Figure S19: **Residue coupling and epistatic effect analysis of key MotAB residues.** (A) Coupling scores (CN values) between target residue pairs, with positive coupling (blue) and negative coupling (red) indicated. (B, C) Box plots of calculated mutational effects for MotA (blue) and MotB (red) residues, using (B) the independent model and (C) the epistatic model. (D) Scatter plot showing the correlation between independent and epistatic model predictions. (E) Distribution of effect differences (epistatic minus independent model), highlighting the magnitude of epistatic interactions.

#### 12 p$K_a$ shifts of Asp in Vacuum Environment

To quantify the extent to which a hydrophobic environment influences proton release and the importance of solvation for this process, vacuum-phase free energy perturbation (FEP) calculations were performed on an isolated aspartic acid (Asp) residue with neutral terminal capping. Furthermore, we found that the free energy difference is substantially elevated, which is 76.32 kcal/mol, resulting in a dramatically large p$K_a$ shift, which is 53.8. This result illustrates that the lack of water molecules will cause an obvious p$K_a$ up shift.

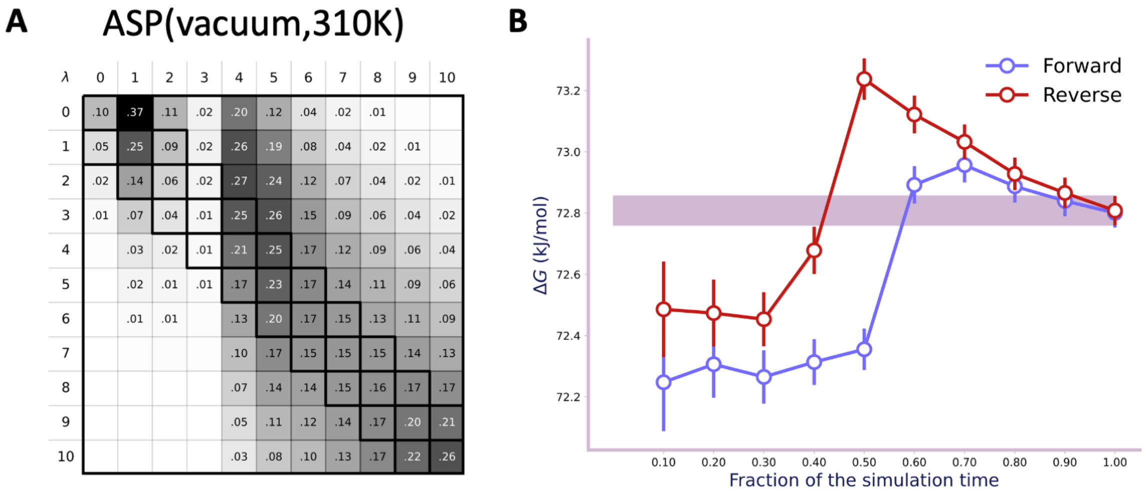


Figure S20: **FEP convergence analysis for aspartic acid in vacuum at 310 K.** (A) Free energy matrix for ASP reference calculation. (B) Forward/reverse FEP convergence plots with converged free energy highlighted.

### 13 Energy of Proton Hydration Solvation Effect

Using the computed pKa values, the deprotonation free energy can be conveniently calculated via the standard reaction free energy equation. Importantly, this free energy contribution can be compensated by the proton hydration (solvation) energy, since the relative free energy of proton hydration falls in the range of -40.21 to -88.07 kcal/mol,[1] corresponding to a strongly exergonic process.

Table S6: The Free Energy of Deprotonation under 300K and pH 7.

| | CpHMD (kcal/mol) | FEP (kcal/mol) | FEP-Vdw (kcal/mol) |
|---|---|---|---|
| $\Delta G_{deprot}^{F-D22}$ | 2.06 | 8.15 | 6.97 |
| $\Delta G_{deprot}^{G-D22}$ | 6.18 | 32.44 | 32.21 |

A

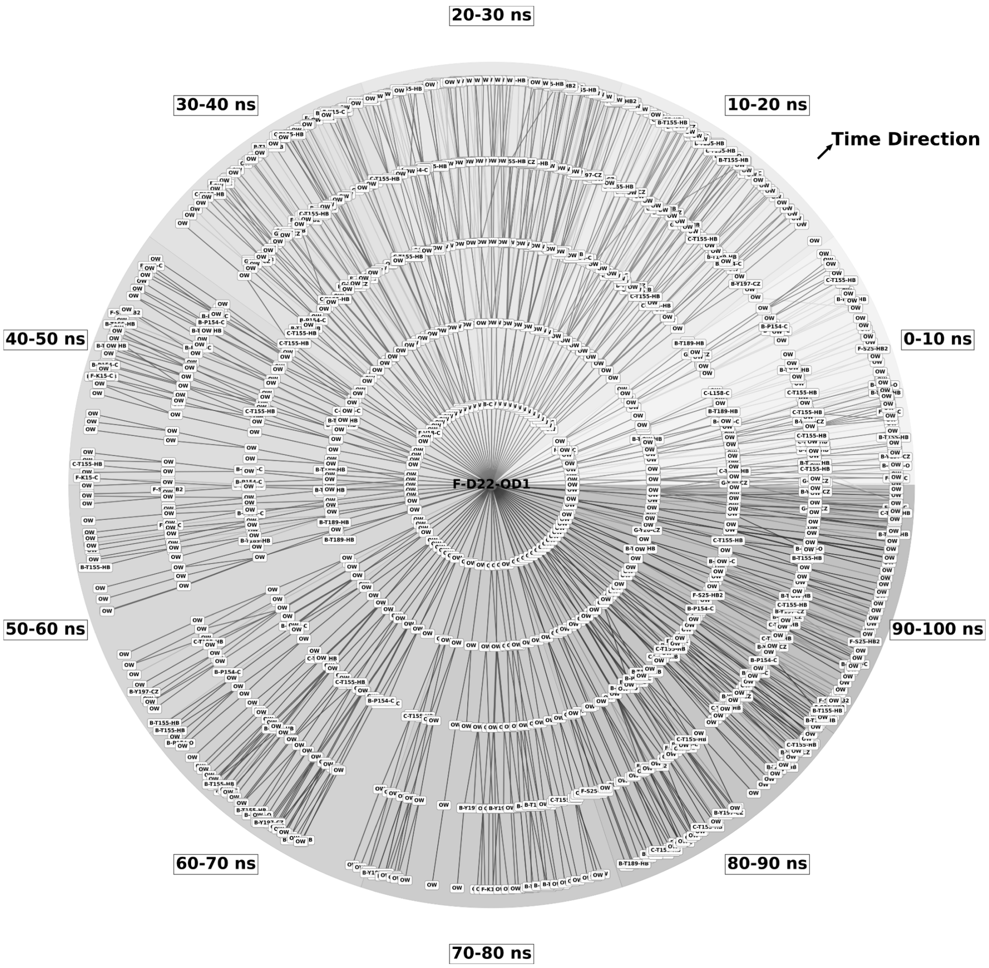


Figure S21: **Time-resolved hydrogen bond network analysis of F-D22-OD1 in the unplugged state from CpHMD simulations.** (A) Radial time-evolution plot visualizing the 5-tier hydrogen bond network originating from the OD1 oxygen atom of F-D22, spanning 0–100 ns of simulation. The concentric rings represent sequential 10 ns time windows, with lines denoting hydrogen bond connections and the gray gradient indicating temporal progression.

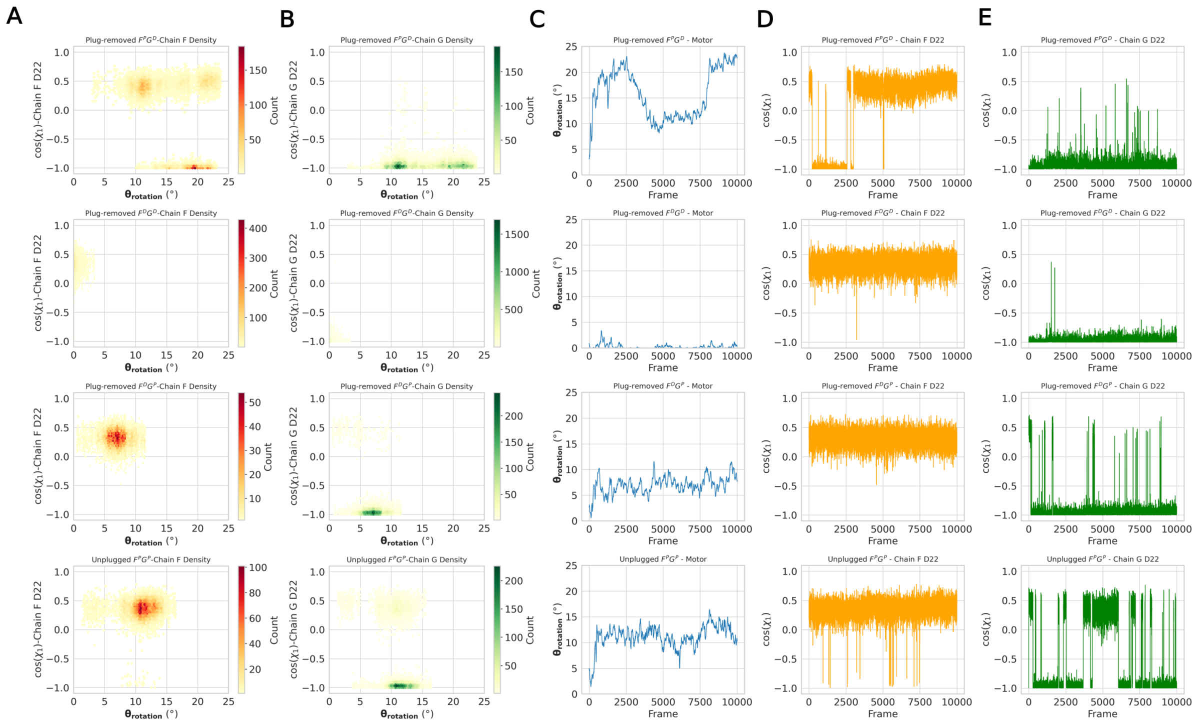


Figure S22: **Conformational coupling between MotA rotation and MotB D22 side-chain dynamics in standard MD simulations.** (A, B) 2D density plots of MotA rotation angle ($\theta_{rotation}$) versus D22 side-chain $\chi_1$ dihedral angle ($\cos(\chi_1)$) across four simulation systems, for (A) F-D22 and (B) G-D22. (C) Time evolution of $\theta_{rotation}$ over 100 ns of simulation for all four systems. (D, E) Time series of $\cos(\chi_1)$ for (D) F-D22 and (E) G-D22, illustrating the dynamic side-chain conformational behavior of D22 residues.

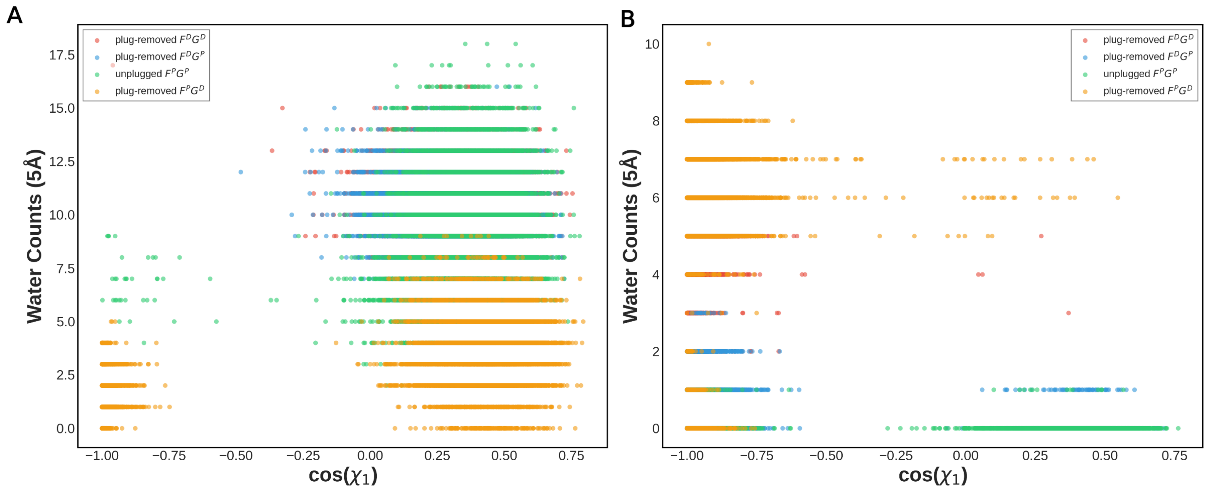


Figure S23: **D22 hydration vs. side-chain $\chi_1$ angle in standard MD simulations.** (A, B) Water counts (5 Å cutoff) plotted against $\cos(\chi_1)$ for (A) F-D22 and (B) G-D22, across multiple protonation/conformational states.

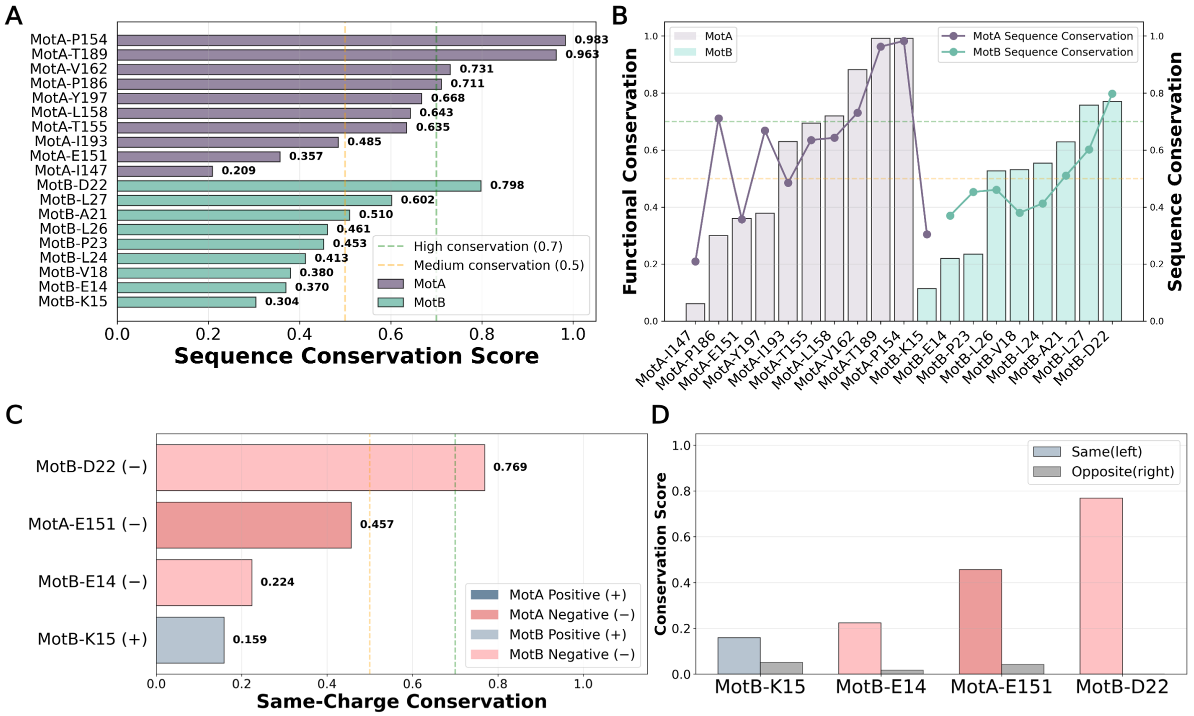


Figure S24: **Sequence and charge conservation analysis of MotAB key residues.** (A) MSA-derived sequence conservation scores for MotA/MotB residues. (B) Sequence vs. functional conservation comparison. (C-D) Same-charge conservation analysis for critical residues, split by charge state and channel orientation.
